# Nutrient signaling, stress response, and interorganelle communication are non-canonical determinants of cell fate

**DOI:** 10.1101/2020.07.17.208603

**Authors:** N Ezgi Wood, Piya Kositangool, Hanaa Hariri, Ashley Marchand, Mike Henne

## Abstract

Isogenic cells can manifest distinct cellular fates for a single stress, however the nongenetic mechanisms driving such fates remain poorly understood. Here, we implement a robust multi-channel live-cell imaging approach to uncover noncanonical factors governing cell fate. We show that in response to acute glucose removal (AGR), budding yeast undergo distinct fates becoming either quiescent or senescent. Senescent cells fail to resume mitotic cycles following glucose replenishment but remain responsive to nutrient stimuli. Whereas quiescent cells manifest starvation-induced adaptation, senescent cells display perturbed endomembrane trafficking and defective nucleus-vacuole junction (NVJ) expansion. Surprisingly, we also show senescence occurs in the absence of lipid droplets. Importantly, we identify the nutrient-sensing linked kinase Rim15 as a key biomarker that predicts cell fates before AGR stress. We propose that isogenic yeast challenged with acute nutrient shortage contain determinants that influence their post-stress fate, and demonstrate that specific nutrient signaling, stress-response, endomembrane trafficking, and inter-organelle tether biomarkers are early indicators for long-term fate outcomes.

## Introduction

To persist in an unpredictable environment, budding yeast cells must adapt to transitions between nutrient rich and poor conditions to maintain viability and proliferative capacity. Such adaptations are achieved through coordinated remodeling of numerous cellular systems encompassing nutrient sensing, stress response, interorganelle communication, metabolic, and energy homeostasis pathways. Depending on the degree and type of nutrient deficiency, budding yeast starvation responses can result in diverse cell fates including quiescence, senescence, pseudohyphal growth, and meiosis (Honigberg, 2016). Importantly, different fates can co-exist in a population of genetically identical cells, implying non-genetic factors also govern cell fate decision-making (Aragon et al., 2008; Argüello-Miranda, Liu, Wood, Kositangool, & Doncic, 2018; De Virgilio, 2012; Laporte, Gouleme, Jimenez, Khemiri, & Sagot, 2018). Although starvation responses in yeast cell cultures have been extensively studied using bulk population-based methods, a key knowledge gap is understanding how the response to nutrient fluctuations is coordinated at the single cell level, and how this response diverges among cells with different fates within a clonal population.

Glucose is a key carbon source for budding yeasts, and its acute removal induces dramatic changes in cell growth and metabolism (Broach, 2012). Glucose starved cells suppress growth through inhibition of conserved TOR and PKA pathways (De Virgilio, 2012), halt mitotic cycles and enter a non-proliferative state marked with significant transcriptional and metabolic remodeling as well as activation of stress responses (Görner et al., 2002). Glucose deprivation also drives organelle remodeling and promotes inter-organelle communication, as evidenced by the expansion of the nucleus-vacuole junction (NVJ), an interorganelle contact site between the nuclear surface and yeast vacuole that physically grows following glucose removal (Hariri et al., 2018; Kvam & Goldfarb, 2006). Remarkably, upon re-exposure to glucose-rich medium, genetically identical yeast cells manifest distinct responses: some cells proliferate and resume budding (defined as quiescent cells) while others do not resume budding despite being metabolically active (defined as senescent cells) (Aragon et al., 2008; De Virgilio, 2012; Laporte et al., 2018; Werner-Washburne, Roy, & Davidson, 2011). How this cell fate is controlled in an isogenic population at the single-cell level is unresolved. Furthermore, how metabolic pathways and organelles communicate to drive these distinct fates remains poorly described. This is particularly true for senescent cells, which display increased lipid storage and lipid droplet (LD) biogenesis, although it is unclear the causative role LDs have in senescence itself (Flor, Wolfgeher, Wu, & Kron, 2017).

Dissecting cell fate within isogenic populations has several technical challenges. First, widely adopted population-based methods often fail, since senescent and quiescent cells are present together. Second, there is no biomarker that unequivocally separates quiescent and senescent cells, apart from their ability to proliferate upon nutrient replenishment (Sagot & Laporte, 2019). This after-the-fact identification necessitates following single cells through the time-course of starvation and nutrient replenishment. Additionally, since multiple pathways take part in adaptive metabolic remodeling, simultaneous measurements of several key pathways are required at the single-cell level to dissect how these pathways contribute to distinct cell fates.

Here we overcome these challenges using a time-lapse imaging platform coupled to a microfluidics device and a computational analysis pipeline (Argüello-Miranda et al., 2018; Doncic, Eser, Atay, & Skotheim, 2013; Wood & Doncic, 2019) that allows us to track single cells before, during, and after acute glucose removal (AGR), while simultaneously following five endogenously-tagged fluorescent biomarkers. Using this system, we simultaneously tracked nutrient signaling, cell cycle, stress, metabolic, and interorganelle contact site markers, dissecting key cellular determinants for response to nutrient availability fluctuations. We show that all cells halt or delay their cell cycle upon AGR regardless of the cell cycle stage they are in. Surprisingly, senescent cells activate stress responses upon AGR, but achieve a distinct cellular response compared to quiescent cells. In particular, we show that senescent cells display altered interorganelle crosstalk as evidenced by their inability to expand their NVJ contacts. However, senescent cells remain responsive to environmental cues throughout and after AGR stress, yet fail to proliferate upon glucose replenishment. In line with this, both senescent and quiescent cells respond to nutrient deprivation by up-regulating lipid storage pathways in the form of lipid droplets (LDs). Surprisingly, we show that LDs are not required for fate differentiation nor senescence. We also demonstrate that Rim15-dependent nutrient sensing can predispose cells toward a particular fate prior to AGR stress. Finally, we apply our biomarker profiles to accurately predict the fates of individual cells during AGR hours before their cell fates manifest, using the Bayesian method of statistical evidence. Collectively, our results indicate that cellular adaptive responses can be pre-disposed to particular fates well before the manifestation of a cell fate.

## Results

### Isogenic cells respond differentially to AGR stress

To interrogate the response of individual cells to acute glucose fluctuations, we developed an assay that merges time-lapse microscopy with microfluidics and allows us to follow individual cells before, during, and after acute glucose removal (AGR) (Video 1). Specifically, yeast in the exponential growth phase were loaded into the microfluidics device and grown for 2hrs in rich media (SCD), and then exposed to AGR (SC) for 10hrs. As expected, upon AGR exposure all cells halted cell growth and budding. Next, cells were re-introduced to glucose rich medium, defined as glucose replenishment, for 4hrs. Remarkably, upon glucose replenishment, some cells resumed mitotic cycles and budding, while others did not (Figure 1A). Following the nomenclature in (Laporte et al., 2018), we denote the growth-resuming cell population as quiescent, and non-resuming population as senescent. Furthermore, time-lapse imaging throughout this process allowed us to distinguish two sub-populations of quiescent cells. As described in detail below, one sub-population completed one cell cycle during the AGR phase. We denote this sub-population as quiescent-committed cells (Figure 1B). The other quiescent sub-population arrested their cell cycle in AGR until glucose replenishment, regardless of the stage of the cell-cycle they were in (Figure 1C). We define these as quiescent-arrested cells, since they arrest their cell cycle during the AGR phase.

**Figure 1.**
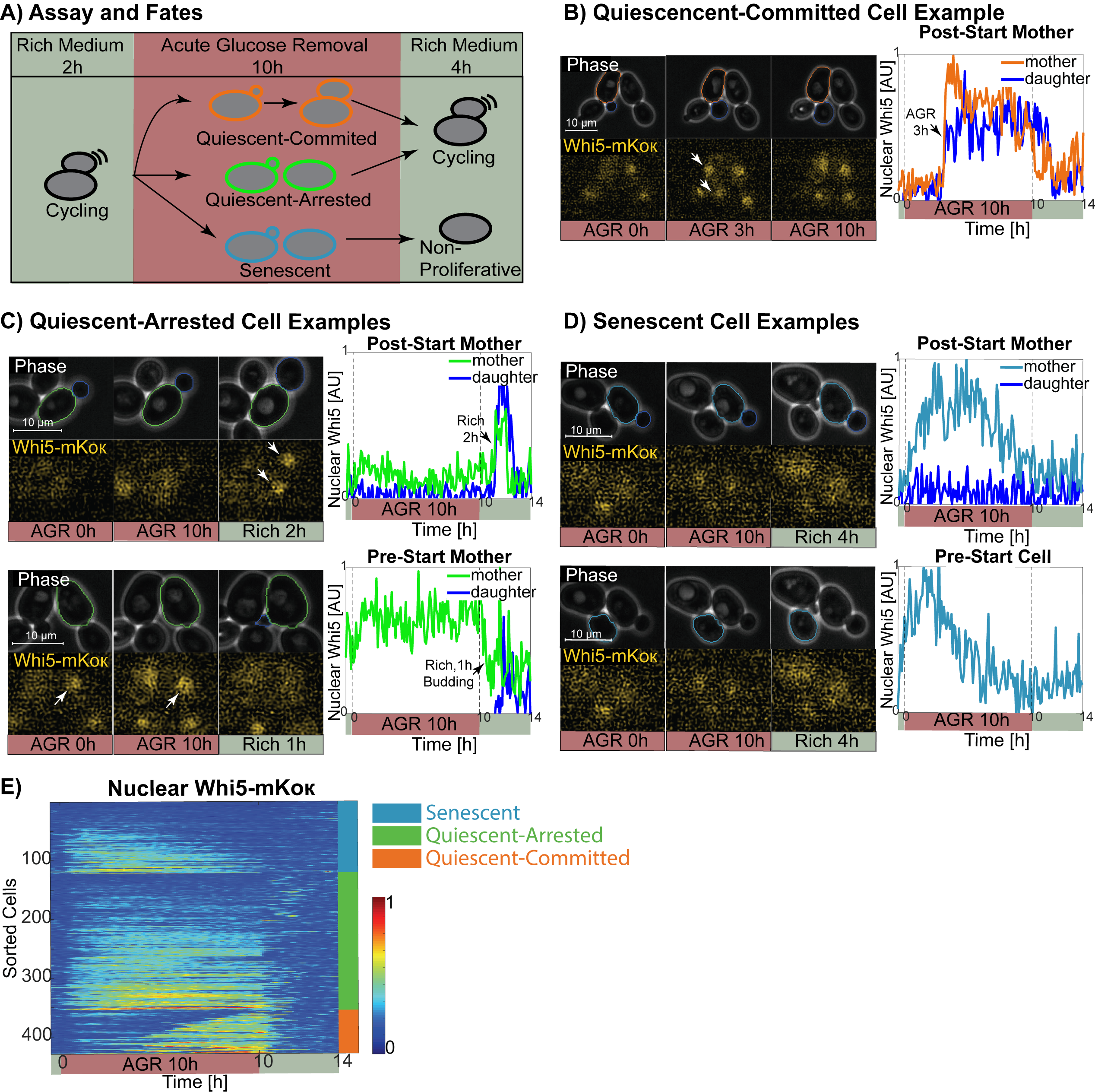
Isogenic cells respond differentially to acute glucose removal (AGR). **A)** Overview of the assay and the fates. Cells of all fates are cycling before AGR. During AGR, quiescent-committed cells complete one cell cycle (orange), whereas quiescent-arrested (green) and senescent (blue) cells halt their cell cycle progression regardless of the cell cycle stage they are in. Upon glucose replenishment, quiescent cells resume mitotic cycling, whereas senescent cells become non-proliferative. **B)** Example quiescent-committed cell. The mother cell has a small bud at the beginning of the AGR (mother-orange, daughter-dark blue, AGR 0h). During AGR, the bud grows, and the cell cycle is completed as seen by the simultaneous translocation of Whi5-mKoκ to mother’s and daughter’s nuclei. After glucose replenishment, the mother and the daughter resume cell cycles. **C)** Example quiescent-arrested cells. Top, the post-start mother (green) arrested with a bud. Bottom, pre-start cell (green). Both cells resume cell cycles after glucose replenishment. **D)** Example senescent cells. The senescent cells do not resume their mitotic cycling, as seen from phase images and the lack of Whi5-mKoκ cycling in and out of the nucleus under replenishment. **E)** Nuclear Whi5-mKoκ traces for WT-MSN2 and WT-RTG1 strains. Cells are first ordered by their cell fate, and then ordered according to their Whi5-mKoκ traces during AGR.

We tracked and segmented individual cells employing algorithms developed in (Doncic et al., 2013; Wood & Doncic, 2019). Cell fates were scored semi-manually using both phase-contrast imaging and fluorescence imaging of the cell cycle marker Whi5-mKoκ, which is an inhibitor of G1-S transition and resides in the nucleus during the G1 phase (Figure 1B-D) (Di Talia, Skotheim, Bean, Siggia, & Cross, 2007). Specifically, we scored quiescence and senescence cell fates by monitoring bud emergence or bud growth after glucose replenishment using phase-contrast imaging. Quiescent-committed fate was scored using both bud growth and Whi5-mKoκ translocation into the nucleus of both mother and daughter during AGR. For budding yeast, the time of commitment to the mitotic cell cycle is denoted as *Start*. Once cells pass Start, they begin DNA replication and the bud concomitantly emerges (Cross, 1995). Thus, we used yeast buds to determine the post-Start cells as in (Jorgensen, Nishikawa, Breitkreutz, & Tyers, 2002).

Budding yeast responses to nutrient deprivation are achieved through coordinated remodeling of numerous cellular systems. To understand the function and interplay of multiple pathways in this cellular decision process, multiple signals must be simultaneously measured at the individual cell level. To this end we utilized our previously developed fluorescent protein tagging and imaging system, which allows for simultaneous monitoring of up to six fluorescent channels (Argüello-Miranda et al., 2018), to monitor protein biomarkers from a diverse array of cellular pathways (Videos 2,3). Here, we focused on biomarkers whose sub-cellular localizations and expression levels reported distinct aspects of cell and organelle metabolism and stress response, including a cell cycle marker (Whi5), a stress-activated transcriptional activator linked to glucose deprivation (Msn2), a transcription factor that coordinates cross-talk between the nucleus and mitochondria to drive respiration (Rtg1), a marker for the nucleus-vacuole junction (NVJ) interorganelle contact site, which expands during glucose starvation (Nvj1), an energy and lipid storage marker that localizes to lipid droplets (LDs) (Erg6), a vacuole homeostasis marker (Vma1), and a stress-induced protein kinase linked to nutrient availability (Rim15) (Figure S1A-C). All lines contained Whi5-mKoκ, Erg6-mTFP1, Vma1-mNeptune2.5, and Nvj1-mRuby3, but differed in their markers for Msn2-mNeonGreen, Rtg1-mNeonGreen, or Rim15-mNeonGreen, and are hence denoted WT-MSN2, WT-RTG1, and WT-RIM15. When reporting joint markers, we pooled the data from WT-MSN2 and WT-RTG1, but presented their individual data in the supplementary. Control experiments were conducted to confirm these fluorescent markers did not grossly affect cell fitness, and to confirm that they could be imaged together with minimal fluorescence bleed-through (Star Methods, Table S1).

### AGR induces cell cycle delay or arrest at any stage of the cell cycle

First, we asked whether a distinct cell fate correlates with the cell cycle stage the cell is in at the initiation of AGR. Because cells are not synchronized at the start of our assay, they were exposed to AGR at an arbitrary stage of their cell cycle. Following AGR initiation, all cells initially halted or postponed their cell cycle regardless of the cell cycle stage they were in, suggesting that even cells in post-Start phase exhibit metabolic control on cell cycle progression (Figure 1B-D). As such, quiescent-committed cells initially halted cell cycle progression, but then completed one cell cycle during AGR and then arrested until glucose replenishment, as evidenced by Whi5-mKoκ nuclear translocation and bud growth during AGR (Figure 1B). We examined the proportion of quiescent-committed cells among all monitored cells, and found they represented 17.5% of cells (Table S2). In line with this, almost all quiescent-committed cells were post-Start (90.7% (68/75), Table S3).

In contrast, quiescent-arrested cells halted their cell cycle during AGR until glucose replenishment, as evidenced by the lack of bud growth and emergence (Figure 1C). Quiescent-arrested cells constituted roughly half of all cells monitored (54.1%, 232/429, Table S2). Of these, a slim majority (64.7%) were pre-Start cells (Table S2). Thus, our data suggests that post-Start cells are more likely to complete their cell cycle during AGR and become quiescent-arrested (p<0.01, Chi-square Test). However, when we pool the quiescent-committed and - arrested cells together, we find that about half of all quiescent cells are pre-Start (51.1%, Table S3), whereas 63.1% of senescent cells are pre-Start (Figure 1D, Table S3). This indicates that the ability to resume mitotic cycles upon glucose replenishment does not depend on the cell cycle stage at the time of AGR initiation (p=0.02, Chi-square test).

Collectively, we find that AGR stress halts or delays the cell cycle of all cells regardless of their cell cycle stage. Quiescent cells which were post-Start upon AGR initiation were more likely to complete their cell cycle during the AGR phase, but the ability to resume mitotic cycles following glucose replenishment is not related to the cell cycle stage during AGR.

### Cell populations display temporally unique Whi5 translocation profiles

To more thoroughly dissect the cell cycle behavior of individual cells relative to one another, we plotted the nuclear Whi5-mKoκ signal of all monitored cells as a heatmap over the 16hrs experimental time-course (Figure 1E, Figure S1F). Although there was a degree of cell-to-cell variation, the nuclear retention of Whi5-mKoκ throughout AGR was distinct for each cell fate population (Figure 1E). For example, quiescent-committed cells completed their cell cycle at different times within the AGR experimental time, upon which Whi5-mKoκ simultaneously entered the nucleus of the mother and daughter cells (Figure1 B,E). Surprisingly, in a sub-population of post-Start quiescent-committed cells, Whi5-mKoκ translocated into the nucleus of the mother during AGR, but then translocated out of nucleus to the cytoplasm, then re-translocated into the nucleus of both the mother and the daughter later in the AGR phase (39.7% (27/68), Figure S1D). Nearly all (85.4%, 70/82)) post-Start quiescent-arrested cells also translocated Whi5-mKoκ into the nucleus during AGR (Figure 1D). Among a majority of quiescent cells (i.e. committed and arrested), nuclear Whi5-mKoκ was generally persistent, remaining until the end of AGR phase (Figure 1E). Collectively, these data suggest that quiescent cells accumulate Whi5-mKoκ in their nucleus during the AGR phase, but surprisingly even without completion of mitotic division Whi5-mKoκ can translocate into the mother cell nucleus during AGR stress.

In contrast to quiescent cells, the majority of senescent cells accumulated nuclear Whi5-mKoκ signal shortly after AGR initiation, but the Whi5-mKoκ nuclear signal gradually declined during the AGR phase (Figure 1E). In line with this, by the end of AGR phase 87.7% of senescent cells lost detectable Whi5-mKoκ nuclear signal. Collectively these observations indicate that cell fate populations respond differentially to sustained AGR stress and display unique temporal signatures of the cell-cycle inhibitor Whi5. It also suggests that Whi5-mKoκ accumulation and retention time in the nucleus during AGR may be a feature that distinguishes cell populations.

### Quiescent and senescent cells respond to AGR with unique stress response signature profiles

When examining senescent cells, we first verified that senescent cell fate was not due to fitness defects originating from their cell cycle phase at AGR initiation, replicative age, doubling time, cell death, or delay in cell cycle resumption (Star Methods). To better characterize senescent cell behavior, we investigated whether senescent cells could sense nutrient stress and whether general stress pathways exhibit unique signature behaviors that correlated with specific cell fates. We generated chromosomally fluorescently-tagged strains of Msn2 or Rtg1 expressed by their endogenous promoters. Msn2 is a stress-responsive transcriptional factor that translocates into the nucleus in response to a variety of stress conditions (Martinez□Pastor et al., 1996; Schmitt & McEntee, 1996). Rtg1 is a transcription factor that enters the nucleus following nutrient starvation and activates genes important to mitochondrial respiration (Butow & Avadhani, 2004; Chen, Sutter, Shi, & Tu, 2017; Liao & Butow, 1993). Remarkably, all cells translocated both Msn2 and Rtg1 into the nucleus within 6-18 min of AGR induction, indicating that cells of all fates sense and respond to AGR stress (Figure 2A,B). However, individual cells within each population displayed different temporal signatures of Msn2 or Rtg1 nuclear export during the AGR phase. All quiescent-committed cells exported Msn2 and Rtg1 out of the nucleus before they completed their cell cycles (Figure 2A,B, Figure S2A,B), suggesting that cell-cycle progression may occur after the downregulation of Msn2 and Rtg1-associated nuclear signaling. In contrast, quiescent-arrested and senescent cells generally exhibited more sustained nuclear Msn2 and Rtg1 signals. In fact, 40.0% (52/130) of the quiescent-arrested cells and 45.8% (33/72) of the senescent cells exhibited extended Msn2 nuclear signal as calculated by comparing the Msn2 nuclear levels between 6^th^ and 7^th^ hrs of AGR to basal Msn2 levels (Figure 2A, Star Methods). By a similar calculation, 11.8% (12/102) of quiescent-arrested cells and 18% (9/50) of senescent cells exhibited sustained Rtg1 nuclear signal (Figure 2B, Star Methods). Collectively, this indicates that different cell populations display temporally distinct stress response nuclear profiles during AGR phase, with quiescent-arrested and senescent cells generally exhibiting more sustained nuclear Msn2 and Rtg1 nuclear signals compared to quiescent-committed cells.

**Figure 2.**
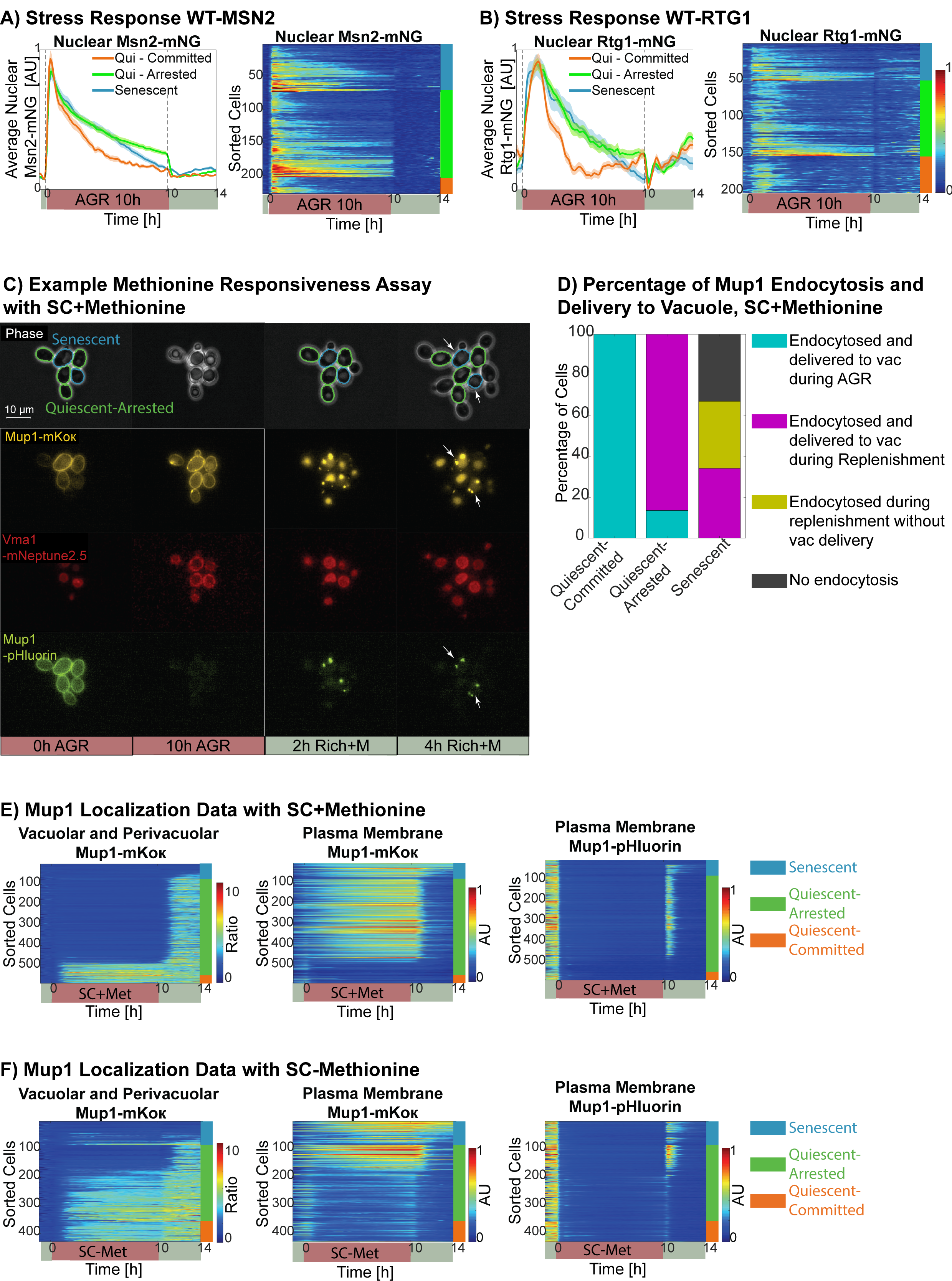
Senescent cells respond to nutrient stress and environmental cues. **A, B)** Average nuclear Msn2-/Rtg1-mNeonGreen traces (mean±SEM) and the heatmap exhibiting individual cell traces. Note that all cells respond to glucose removal by translocating Msn2/Rtg1 to their nuclei. **C)** Example Methionine Responsiveness Assay. Cells grown in media lacking methionine (SCD-Met) are loaded to the microfluidics device and exposed to AGR with SC media containing methionine (SC+Met). Senescent cells are marked blue and quiescent-arrested cells are marked green in the phase images. Note that although the senescent cells endocytose Mup1 as seen by the punctae in Mup1-mKoκ and Mup1-pHluorin channels, they do not successfully carry them to the vacuole evidenced by the lack of quenching of the Mup1-pHluorin signal (the punctae are pointed with white arrows, compare to quiescent cells with quenched signals). Vma1-mNeptune2.5 marks the vacuole. This colony is also shown in Video 4. **D)** Percentage of cells that endocytosed Mup1 and delivered it to the vacuole by cell fate during methionine responsiveness assay with SC+Met. vac:vacuole. **E, F)** Mup1 localization quantification for methionine responsiveness assay with SC+/-methionine. In the first phase of the assay, cells are in SCD-Methionine, and thus Mup1-mKok/pHluorin is localized on the cell membrane as seen by the lack of vacuolar/perivacuolar Mup1-mKoκ signal, and the presence of plasma membrane Mup1- mKoκ and Mup1-pHluorin signals. Upon glucose removal, some cells endocytose Mup1 and carry it into the vacuole as seen by their vacuolar/perivacuolar Mup1-mKoκ signal, and the lack of plasma membrane Mup1-mKoκ signal. Note that the Mup1-pHluorin signal disappears for every cell during AGR, because for cells which carry Mup1 into the vacuole, pHluorin signal is quenched due to the low pH in the vacuole. For cells that have Mup1 on the membrane, the signal is quenched due to cytoplasmic acidification. For these cells, plasma membrane Mup1-pHluorin signal briefly comes back after glucose replenishment, until they carry Mup1 into the vacuole during replenishment.

### Senescent cells execute endocytosis in response to nutrient cues, but exhibit defects in post-endocytic endomembrane trafficking

Since senescent cells appeared to sense AGR as evidenced by their nuclear translocation of Msn2 and Rtg1, we next tested whether they are capable of coordinated responses to other stimuli. We monitored the endocytic uptake and trafficking of the methionine permease Mup1 as a stimuli-responsive test (Video 4-6). Mup1 is the primary methionine permease of budding yeast. When cells are grown in nutrient-rich media lacking methionine, Mup1 is primarily located on the plasma membrane (PM). Upon exogenous methionine addition, Mup1 is endocytosed and trafficked into the vacuole for degradation via ESCRT-dependent protein sorting. Similarly, if cells are glucose starved, Mup1 and other surface permeases are bulk endocytosed and delivered into the vacuole lumen to be broken down and supply the cell with amino acids to survive (Lang et al., 2014). As such, Mup1 trafficking and sub-cellular localization enables us to monitor a cell’s responsiveness to nutrient deprivation.

We generated a diploid yeast strain where one chromosomal copy of Mup1 was tagged with the pH-sensitive fluorophore pHluorin, whose fluorescent signal is quenched when the cytoplasm is acidified upon glucose deprivation, or when it is delivered into the vacuole acidic lumen (William Mike Henne, Buchkovich, Zhao, & Emr, 2012; Lin, MacGurn, Chu, Stefan, & Emr, 2008). The second Mup1 chromosomal copy was tagged with the pH-resistant fluorophore mKoκ. We also labeled the vacuole ATPase subunit Vma1 with mNeptune2.5 so that Mup1-mKoκ vacuole delivery could be directly monitored (Graham, Flannery, & Stevens, 2003).

First, we monitored Mup1-pHluorin and Mup1-mKoκ localizations in yeast grown in rich media lacking methionine (SCD-Met), allowing Mup1 to accumulate at the PM. Cells were then exposed to AGR in media either containing or lacking methionine (SC+Met, SC-Met) for 10hrs, followed by glucose replenishment in media containing methionine (SCD+Met), which should stimulate any residual Mup1 uptake via endocytosis (Lin et al., 2008). During AGR in either SC+Met and SC-Met media, all quiescent-committed cells endocytosed Mup1-mKoκ off the PM before they completed their cell cycles (Figure 2D-F Figure S2C, Video 6). Following its uptake, Mup1-mKoκ was also observed in the vacuole lumen of quiescent-committed cells, indicating quiescent-committed cells can successfully complete Mup1 endocytosis and vacuole delivery. In contrast, 86.4% (408/472) of quiescent-arrested cells challenged with AGR in SC+Met media retained Mup1-mKoκ at the PM, indicating that glucose starvation was generally not sufficient to induce Mup1-mKoκ endocytosis in this sub-population (Figure 2D,E). However, in AGR lacking both glucose and methionine (SC-Met), 71.8% (196/273) of quiescent-arrested cells endocytosed Mup1-mKoκ and trafficked it into the vacuole lumen (Figure 2F, Figure S2C). Collectively, this data indicates that during AGR, most quiescent cells can endocytose Mup1-mKoκ, but quiescent-arrested cells preferentially retain Mup1-mKoκ at the PM if there is methionine in the culture medium.

In contrast, senescent cells displayed different Mup1 trafficking profiles than quiescent cells. During the AGR phase in SC+Met media, Mup1-mKoκ remained at the PM and was not endocytosed (Figure 2D,E). However, upon glucose replenishment with SCD+Met, 67% (51/76) of the senescent cells endocytosed Mup1-mKoκ, whereas 32.9% (25/76) retained Mup1-mKoκ at the PM. This indicates that although senescent cells are non-proliferative, a majority are responsive to environmental cues and can execute an endocytic response following sustained AGR stress (Figure 2C-E). In line with this, 47.0% (39/83) of senescent cells internalized Mup1-mKoκ off the PM during AGR phase when cultured in SC-Met media, and 28.9% (24/83) of senescent cells endocytosed Mup1-mKoκ during the subsequent glucose replenishment (Figure 2F, Figure S2C).

Next, we focused on Mup1-pHluorin to monitor the acidification of the cytoplasm, a known consequence of glucose starvation in yeast (Dechant et al., 2010; Joyner et al., 2016). Immediately following AGR exposure the cytoplasm of all cell populations acidified as evidenced by the quenching of Mup1-pHluorin signal at the PM (Figure 2E,F). As expected, glucose replenishment returned the cytoplasm back to a neutral pH, allowing us to examine the sub-cellular localization of Mup1-pHluorin. This revealed that although a majority of senescent cells were able to successfully internalize Mup1 upon glucose replenishment, Mup1-pHluorin was not always successfully delivered into the vacuole lumen, where the pHluorin signal is normally quenched. In fact, in 49% (25/51) of senescent cells that endocytosed Mup1 after SC+Met AGR treatment, Mup1-pHluorin puncta accumulated in peri-vacuolar punctae within the cytoplasm (Figure 2C, blue lined cells and white arrows). Consistent with this, Mup1-mKoκ also accumulated as bright punctae in the peri-vacuolar space of these cells, indicating a defect in vacuolar sorting of Mup1 (Figure 2C). In contrast, all quiescent cells were able to eventually deliver and quench Mup1-pHuorin into the vacuole as evidenced by its fluorescence loss. Indeed, Mup1-mKoκ could be observed in the vacuole lumen of quiescent cells following glucose replenishment (Figure 2C, green lined cells, Figure 2D-F). This suggests that although senescent cells can execute Mup1 endocytosis following AGR, some stage of post-endocytic endosomal trafficking and/or vacuole delivery is compromised. In contrast, both quiescent sub-populations appear to successfully deliver Mup1 into the vacuole lumen following endocytosis, as evidenced by increased vacuolar Mup1-mKoκ signal and quenched intracellular Mup1-pHluorin signal (Figure 2E,F).

In summary, in both quiescent and senescent cells the cytoplasm acidifies during AGR stress and they can generally execute endocytosis, indicating they have the ability to remodel their PM in response to stress stimuli. Quiescent cells also successfully deliver Mup1 into the vacuole lumen. In contrast, only about half of senescent cells that endocytose Mup1, successfully delivered Mup1 into the vacuole lumen where Mup1-pHluorin signal is quenched, suggesting defects in endosome maturation or vacuole homeostasis that may underlie defects in senescent cell behavior.

### Relative V-ATPase assembly differs among quiescent and senescent cells

Since senescent cells displayed defects in post-endocytic endomembrane trafficking, we next investigated whether vacuole homeostasis was altered by monitoring the vacuolar ATPase (V-ATPase), the proton pump that regulates the intraluminal vacuole pH (Graham et al., 2003). The yeast V-ATPase is composed of two subcomplexes, whose reversible assembly controls V-ATPase function. Upon glucose removal the V_1_ subcomplex dissociates, whereas the V_0_ subcomplex remains membrane associated. Due to its dissociation, the coefficient of variation (CoV) of V_1_ subcomplex signal intensity has been shown to be a marker for relative V-ATPase assembly (Dechant et al., 2010). Thus, we monitored the CoV of Vma1-mNeptune2.5, a subunit of the membrane-dissociating V_1_ subcomplex. As expected, the normalized Vma1-mNeptune2.5 CoV decreased for all cells during AGR. However, the signal decreased more drastically for senescent cells compared to quiescent cells (Figure S2D). Specifically, the CoV normalized to 1 at the beginning of AGR decreased to 0.65±0.01 or quiescent cells, but decreased to 0.59±0.02 for senescent cells (mean±SEM). Upon glucose replenishment, Vma1-mNeptune2.5 CoV increased for both quiescent and senescent cells, however, it did not recover in senescent cells to the degree observed in quiescent cells (Figure S2C). As a control, we also monitored the normalized CoV of the Vph1-GFP, a subunit of the V_0_ subcomplex that remains membrane bound during AGR. As expected, Vph1-GFP CoV did not significantly change during AGR (Figure S2E, drops to 0.991±0.033 for quiescent and 0.944±0.023 for senescent cells, mean±SEM). Collectively, the more drastic CoV change of Vma1-mNeptune2.5 in senescent cells is consistent with altered vacuole homeostasis in the senescent cell population.

### Quiescent cells enlarge their NVJ contacts during AGR, but senescent cells do not

The vacuole does not function in isolation, and is connected to other organelles including the nucleus that enable it to act in concert with other organelles to respond to stress. Since senescent cells exhibited altered endomembrane trafficking and vacuole homeostasis, we next investigated whether the nucleus-vacuole junction (NVJ), a disc-shaped interorganelle contact site that expands during nutrient deprivation, was also altered (Hariri et al., 2018; W Mike Henne & Hariri, 2018; Pan et al., 2000). We monitored chromosomally tagged Nvj1-mRuby3 to evaluate NVJ expansion during the AGR experiment (Figure 3A, Figure S3A,B). Remarkably, 87.7% of quiescent cells displayed significantly increased Nvj1-mRuby3 signal during AGR, consistent with NVJ expansion. However the opposite was observed for senescent cells; 80.3% of senescent cells failed to increase Nvj1-mRuby3 signal during AGR, indicating a defect in NVJ expansion (Figure 3B,C; Star Methods). This suggests that failure to up-regulate Nvj1 protein and expand the NVJ contact during AGR stress correlates with senescent cell fate.

**Figure 3.**
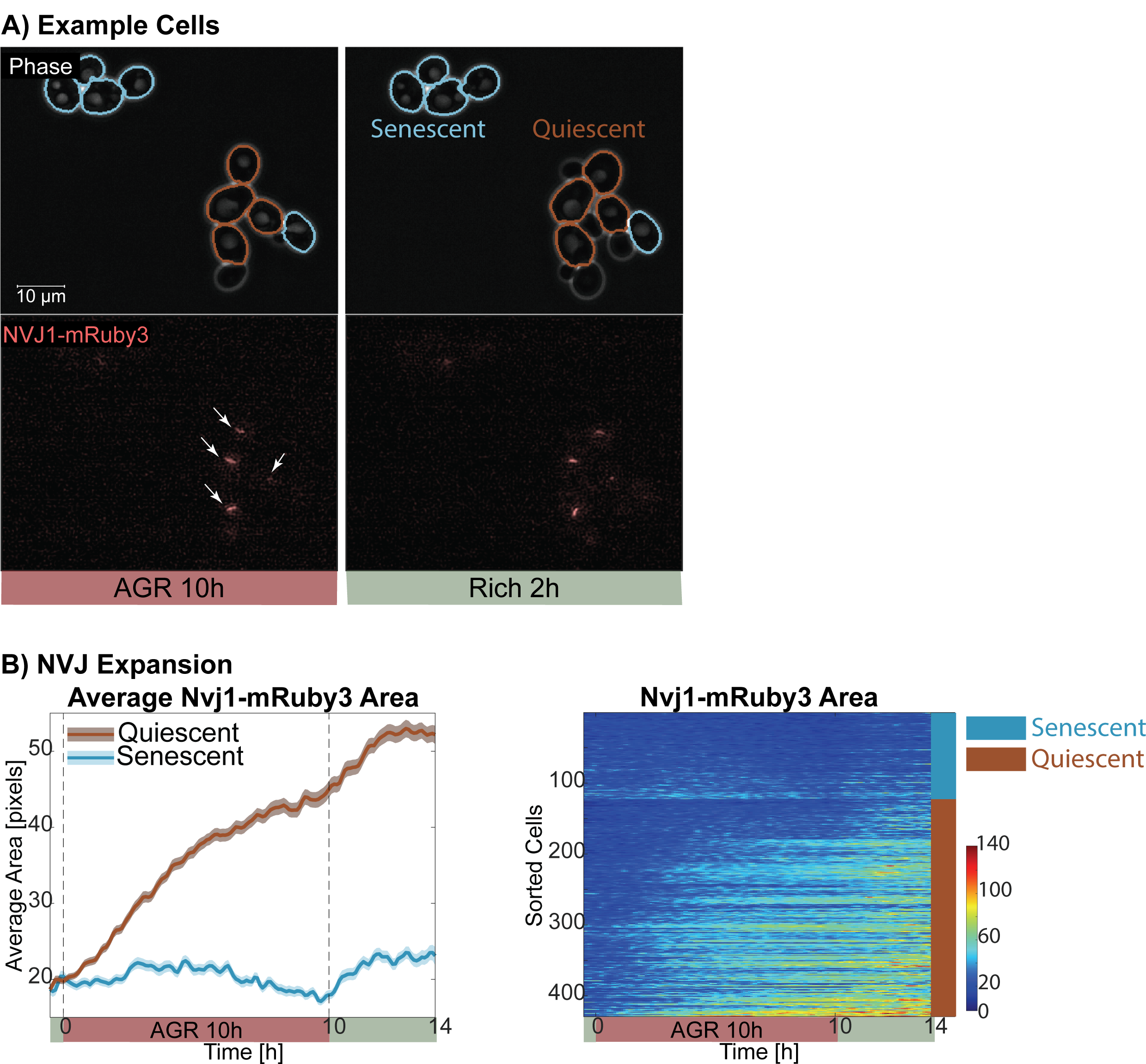
Quiescent cells enlarge their nucleus vacuole junction contacts during AGR, but senescent cells do not. **A)** Example quiescent (brown) and senescent (blue) cells. Note that the Nvj1-mRuby3 expands in quiescent cells (pointed with white arrows), however, it does not expand in senescent cells. At 2h in rich medium, quiescent cells are seen having new buds or growing an existing bud. This colony is also shown in Video 1 and 3. **B**,**C)** NVJ expansion quantification for WT-MSN2 and WT-RTG1: Left, Average Nvj1-mRuby3 area is given for quiescent (brown) and senescent (blue) cells (mean±SEM). Right, individual cell traces are given as a heatmap. Before AGR, cells do not have expanded Nvj1-mRuby3 signal, and our software picks a basal/background area for the Nvj1-signal about 20 pixels.

Next, we queried whether loss of the NVJ via genetic deletion of *NVJ1* would alter cell fates following AGR treatment. We monitored WT and *nvj1*Δ yeast exposed to AGR, followed by glucose replenishment. Remarkably, *nvj1*Δ yeast displayed similar proportions of senescent and quiescent cells compared to WT (Table S2, Chi-square test, p>0.1). We conclude that NVJ expansion is a correlative indictor of senescent cell fate after AGR but is not required for quiescent cell fate. We also cannot exclude the possibility that other ER-vacuole tethers such as Mdm1 or Ltc1/Lam6 may compensate for the loss of Nvj1 (W Mike Henne et al., 2015; Murley et al., 2015).

### Cells exhibit enhanced lipid storage during AGR, but LDs are not required for cell fates

The NVJ expands during glucose starvation, and serves as a platform for the local production of LDs, which store lipids during nutrient deprivation to promote cell survival (Hariri et al., 2018; Radulovic et al., 2013). Once glucose is resupplied LDs can be mobilized to support new membrane synthesis and growth (Kohlwein, Veenhuis, & van der Klei, 2013). Since senescent cells generally failed to increase Nvj1-mRuby3 protein levels, we next investigated whether they also exhibited defects in lipid metabolism and LD biogenesis. To interrogate this, we monitored LD biogenesis by imaging Erg6-mTFP1, an established LD marker and enzyme in the ergosterol biosynthetic pathway (Jacquier et al., 2011; Kristan & Rižner, 2012; Seo et al., 2017). Following AGR, Erg6-mTFP1 signal increased in every cell subpopulation, consistent with elevated LD biogenesis upon glucose removal. The fold increase in the senescent population was significantly lower than the quiescent population (Figure 4A-C, Figure S4A-D). Consistent with this increase in Erg6-mTFP1 signal, yeast cultures exhibited a slight elevation in triglycerides (TG), and a significant elevation in ergosterol-esters (SE) during AGR exposure, as determined by thin layer chromatography (TLC) (Figure 4D). Remarkably, glucose replenishment induced a further increase in TG, but SE levels returned to pre-AGR levels, consistent with LD mobilization of SEs to provide sterols for new membrane synthesis (Figure 4D). In line with this, Erg6-mTFP1 signal decreased on average 30.1% in quiescent cells. In contrast, Erg6-mTFP1 signal only decreased 8% in senescent cells. This Erg6-mTFP1 signal decrease may be due to LD mobilization and/or due to protein dilution in cells which resumed budding.

**Figure 4:**
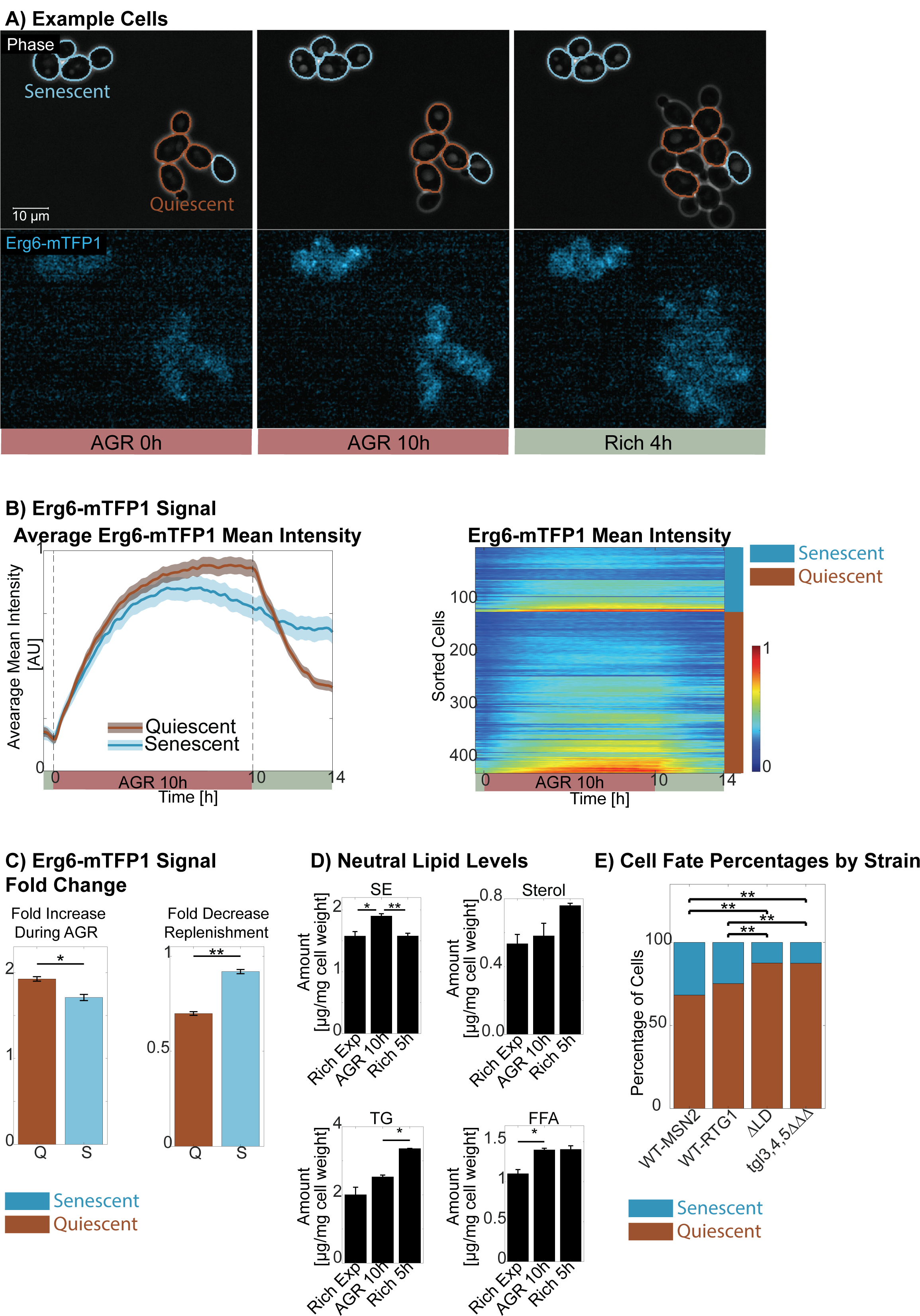
Cells exhibit enhanced lipid storage during AGR. **A)** Example quiescent (brown) and senescent (blue) cells. This colony is also shown in Video 1 and 3. **B)** Erg6-mTFP1 signal quantification: Left, Average Erg6-mTFP1 mean intensity (mean±SEM). Right, individual cell traces for Erg6-mTFP1 mean intensity. **C)** Fold increase during AGR is calculated by determining the ratio of the Erg6-mTFP1 mean intensity at the end of AGR and at the beginning of AGR (1.93±0.03/1.71±0.04 Quiescent /Senescent (mean±SEM)) Fold decrease during replenishment is calculated by determining the ratio of the Erg6-mTFP1 mean intensity at the end of four hour replenishment to the end of AGR (0.70±0.01/0.92±0.01 Quiescent/Senescent (mean±SEM)). Q:Quiescent, S:Senescent. *p<0.05, **p<0.01, Kolmogorov-Smirnov test. **D)** Neutral lipid levels: quantification of the thin layer chromatography experiment. SE:sterol ester, TG: Triacylglycerol, FFA: Free fatty acid.*p<0.05, **p<0.01, *t*-test. **E)** Cell fate percentages by the strain.**p<0.01, Chi-square test.

LDs have been observed to accumulate in senescent mammalian cells, and cell senescence correlates with increased lipid synthesis (Flor et al., 2017; Lizardo, Lin, Gokcumen, & Atilla-Gokcumen, 2017). However, whether LDs are operationally required for cell senescence is unclear. Given that LDs are lipid reservoirs that can contribute to yeast survival during long-term starvation (Seo et al., 2017; Wang, 2015), we next examined whether LDs are required for any cell fates following AGR stress. We compared the fates of WT cells with LD-deficient cells that lacked the two genes required to produce TG (Dga1 and Lro1) and SE (Are1 and Are2) (strain denoted as ΔLD). Remarkably, this genetic perturbation caused a slight decrease in the number of senescent cells, but senescent cells were still detected within the ΔLD strain (Table S2, Figure 4E, Figure S4E). We also examined whether yeast lacking the three major TG lipases also displayed altered cell fate proportions (strain denoted *tgl3,4,5*ΔΔΔ). The *tgl3,4,5*ΔΔΔ yeast closely mimicked the ΔLD yeast with slightly reduced numbers of senescent cells (Table S2, Figure 4E). Collectively, this suggests that cells of all fates increase LD storage during AGR stress. However, surprisingly LDs are not required for quiescent nor senescent cell fates in our experimental framework.

### Pre-AGR Rim15 levels are predictive of post-AGR cell fate

Thus far, our data indicate that quiescent and senescent cells exhibit differences in the profiles of stress response transcription factors, endomembrane trafficking dynamics, and NVJ expansion during or after AGR stress. However, prior to AGR stress both senescent and quiescent cells exhibit similar levels of the protein biomarkers associated with these pathways, indicating that the abundances of these biomarkers provide no predictive power to designate cell fate prior to AGR (Figure 2A,B,E,F, Figure 3B, Figure 4B). Given the possibility that cell fates might be emanating from some pre-existing heterogeneity within the population, we queried whether we could identify a biomarker whose fluorescent signal *before* AGR correlated to distinct cellular behaviors *after* AGR stress.

We hypothesized that proteins involved in nutrient signaling, particularly those that are naturally expressed at low levels in the cell, are strong candidates for factors which may be indicative of cell fate prior to nutrient stress exposure. This is because low abundance proteins display intrinsic cell-to-cell abundance variability, providing a mechanism for isogenic diversity. Using a candidate-based approach, we selected Rim15 as a potential predictor for several reasons. First, Rim15 is an effector kinase that is downstream of at least four major nutrient signaling pathways, and regulates cell proliferation (Smets et al., 2010). Second, Rim15 is normally expressed at low levels in cycling cells (Lawless et al., 2016; Newman et al., 2006). Thus, we fluorescently tagged endogenous Rim15 (Rim15-mNeonGreen) and monitored its mean intracellular intensity throughout our experimental protocol (Figure 5A, Video 7).

**Figure 5:**
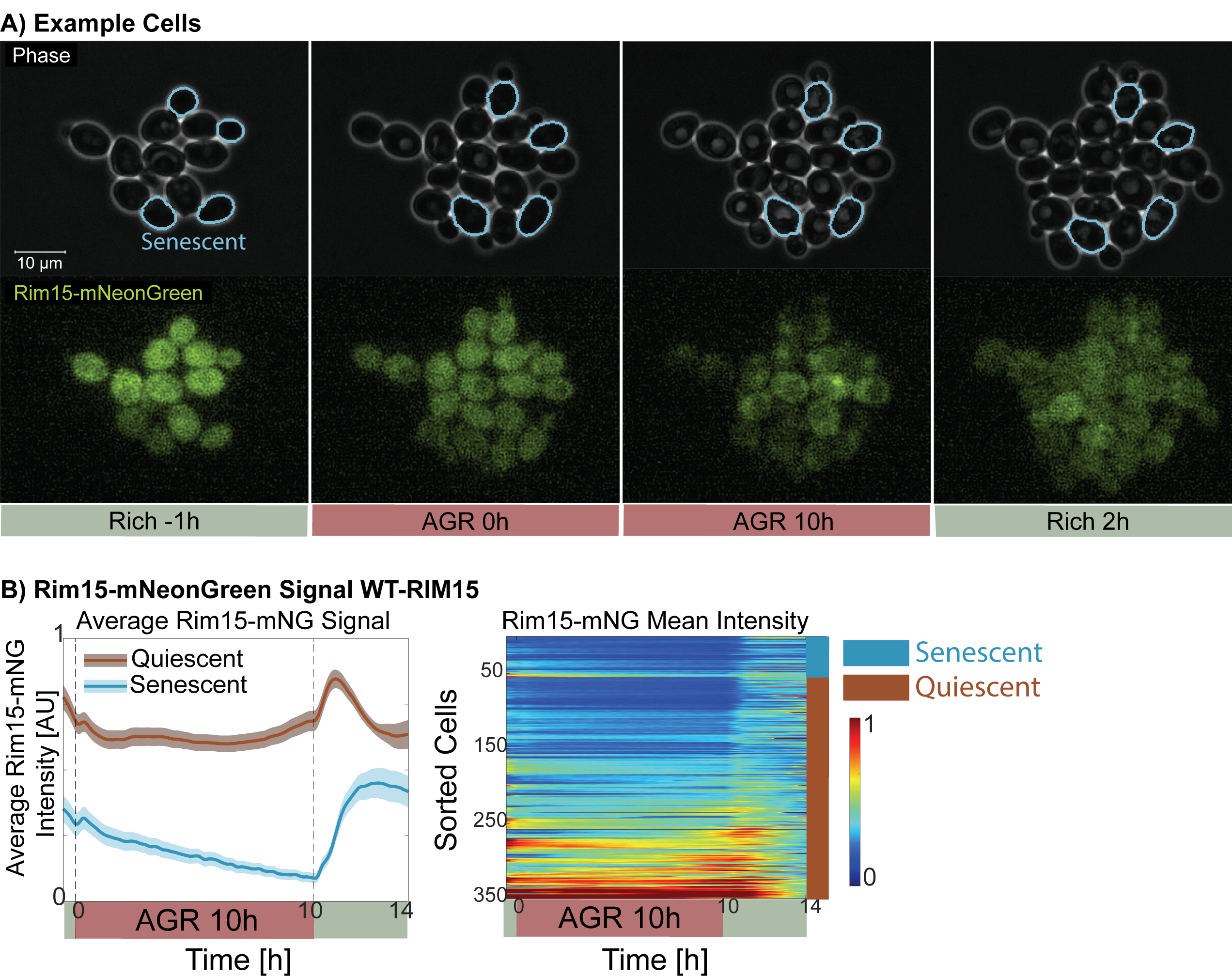
Pre-AGR Rim15 levels are predictive of post-AGR cell fate. **A)** Example quiescent (unmarked) and senescent (blue) cells. This colony is also shown in Video 7. **B)** Rim15-mNeongreen signal quantification: Left, Average (mean±SEM), Right, heatmap of individual cell traces.

Strikingly, elevated Rim15 levels prior to AGR were highly correlated with quiescent cell fate, whereas cells with low Rim15 before AGR were more likely to be senescent (Figure 5B, Figure 5SA). Specifically, 95.9% (47/49) the cells with highest Rim15-mNeongreen levels were quiescent determined as cells that have higher Rim15-mNeonGreen levels than one standard deviation from the mean at the initiation of AGR. In contrast, 88.9% (48/54) of senescent cells had lower than average Rim15-mNeongreen intensity at the beginning of AGR. This percentage is 59.8% (177/296) for quiescent cells. This indicates that Rim15 levels before AGR correlate with cell fate after AGR.

### Cells ‘decide’ their fate prior to glucose replenishment

Next, using the biomarkers studied so far, we queried whether we could predict the fates of individual cells before glucose replenishment, and if so, how long before glucose replenishment. In (Argüello-Miranda et al., 2018), we developed a theoretical framework using statistical evidence (Jaynes, 2003) to convert single cell measurements into cell fate probabilities to predict the fates of individual cells hours before the cell fates are manifested. Briefly, this is approached as a binary hypothesis testing problem, where we compare for a given cell the probabilities to become senescent or quiescent. To convert single cell measurements into probabilities of cell fate, we first pool our measurements from cells of a given fate at a given time to estimate population distributions. Next, using these distributions we calculate the likelihood of a given cell to belong to either of the two populations (Star Methods).

To illustrate this method, and to describe the evolution of population distributions, we used as an example Nvj1-mRuby3 mean intensity, the most powerful single parameter predictor for the fates for our cells (Figure 6A). Note that the Nvj1-mRuby3 signal distributions for quiescent and senescent cells were indistinguishable at the start of AGR, however, they gradually diverged throughout the AGR phase (quiescent (brown) and senescent (blue) distributions, Figure 6A). In fact, this was the case for all the biomarkers interrogated, except for Rim15 (see below). Using Nvj1-mRuby3 distributions, we then predicted the cell fate of each cell and reported the percentage of correctly predicted cells using this approach (Figure 6A, back axis, dark blue). At the beginning of AGR, using Nvj1-mRuby3 data around 50% of the cells are accurately assigned to their correct fate, which is what a random guess would achieve (dark blue line, Figure 6A). However, as the population distributions of Nvj1-mRuby3 diverge, the percentage of correct predictions increases later in the AGR phase. In fact, we accurately predict the fates of 81.9% of cells by the end of AGR and before glucose replenishment.

**Figure 6:**
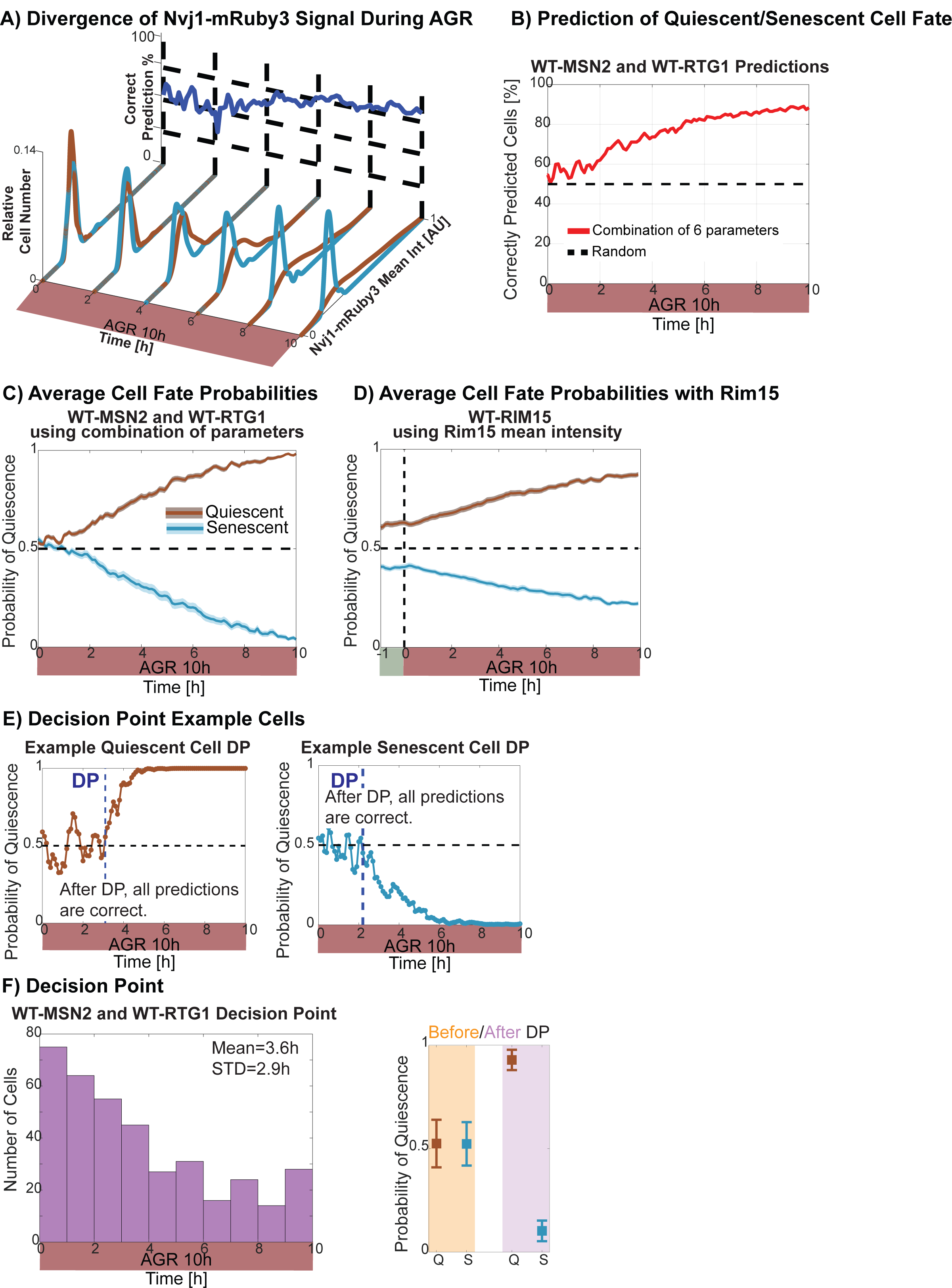
Cells ‘decide’ on their fate before glucose replenishment. **A)** Quiescent (brown) and senescent (blue) Nvj1-mRuby3 mean intensity population distributions throughout the AGR phase and the percentage of correctly predicted cells calculated utilizing these distributions (back axis, dark blue). **B)** Percentage of correctly predicted cells combination of multiple markers (red), WT-MSN2 and WT-RTG1 strains. 50% marks what a random guess would achieve in a binary decision (bold dashed line). See also Figure S6A. **C)** Average quiescence fate probabilities for quiescent (brown) and senescent (blue) cells calculated using the combination of markers from Figure 6B, red line. See also Figure S6C,D. **D)** Average quiescence fate probabilities for quiescent (brown) and senescent (blue) cells calculated using Rim15-mNeonGreen mean intensity. **E)** The evolution of the quiescence fate probability for an example quiescent (left, brown) and for an example senescent cell (right, blue), and the calculation of the Decision Point (DP). **F)** Left, The histogram of DP times and average probability of quiescence before and after DP (mean±SEM), WT-MSN2 and WT-RTG1 strains.

Next, we combined the population distribution information from multiple biomarkers used in this study. Using this multi-parameter approach, by the completion of AGR phase we could correctly predict the fates of 88.3% of all cells (Figure 6B). Specifically, for WT-MSN2 we combined information from Nvj1, Msn2, Whi5, and Vma1 biomarkers, whereas for WT-RTG1 we combined information from Nvj1, Rtg1, Whi5, and Vma1 biomarkers (Figure S6A). Note that since Erg6-mTFP1 signal was increasing in both populations during AGR, it was not a strong predictor of cell fate. For a detailed discussion of parameters and parameter selection methodology see Star Methods. Notably, cell size was also not a good predictor for quiescence/senescence decision (Figure S6B), although it is a strong predictor of meiosis with over 90% accuracy (Argüello-Miranda et al., 2018; Day et al., 2004). This result is consistent with (Laporte et al., 2018), where it is shown that cell volume does not effect a cell’s propensity for quiescence.

The fact that we accurately predicted the fates of 82.1-89.0% of the cells in the last four hours of AGR (Figure 6B) indicates that quiescent and senescent populations exhibit already diverged cellular biomarkers hours before glucose replenishment. Collectively, our data suggest that within our experimental system, yeast cells “decide” a fate (i.e. become predisposed to a specific fate), hours before the manifestation of that cell fate (to cycle or not).

### Rim15 abundance contains information about cell fates before AGR

Next, we asked when individual cells ‘decide’ on their fate, and thus we switched our focus from the percentage of correctly predicted cells to dissecting an individual cell’s fate probabilities over time. Specifically, we examined the evolution of quiescence cell fate probabilities throughout the AGR phase using cells whose fates were correctly predicted at the end of the AGR phase utilizing the combination of markers used in Figure 6B (Figure 6C, Figure S6C, average probability of quiescence for quiescent (brown) and senescent (blue) cells, mean±SEM). Note that the probability of quiescence was around 0.5 for both quiescent and senescent cells at the beginning of AGR (0.53±0.01 / 0.55±0.02 for Quiescent/Senescent, mean±SEM). However, this probability steadily increased for quiescent cells during AGR phase and decreased for senescent cells and became 0.98±0.004 / 0.03±0.008 for Quiescent/Senescent cells at the end of AGR (mean±SEM).

One exception to this trend was the biomarker Rim15-mNeonGreen. Whereas both single parameters and the combination of parameters gave the probability of quiescence near 0.5 at the beginning of AGR (Figure 6C, Figure S6C,D), the probability of quiescence calculated using Rim15-mNeonGreen is 0.63±0.01 for quiescent cells (N=206) and 0.41±0.01 (N=47) for senescent cells (mean±SEM) at the start of AGR phase (Figure 6D). In fact, even one hour before AGR initiation these probabilities were 0.61±0.01 and 0.41±0.01 respectively for quiescent and senescent cells. Thus, Rim15-mNeonGreen is the only biomarker that appeared to encode information about cell fate even *prior* to AGR. Although by itself Rim15-mNeongreen signal was not a strong predictor for all cells (compare Figure S6E to Figure 6B and S6A), when we restricted our predictions to cells with signal one standard deviation above or below the average Rim15-mNeonGreen signal, we accurately predicted their at 88.1%±3.4% (mean±std) throughout the entire AGR phase (Figure S6F). Applying the same procedure for predictions with Nvj1-mRuby3 area and mean intensity did not improve their overall predictive power (Figure S6F).

### The cell fate decision occurs early in AGR phase

Since multiple biomarkers significantly diverge among cells of different fates throughout AGR, and the combination of multiple markers provides both a higher percentage of correctly predicted cells at the end of AGR, as well as more confidence in these predictions as reflected by the cell fate probabilities, we conducted further analysis with biomarker combinations. When we analyzed the cell fate probabilities of individual cells calculated using the combination of parameters, we observed that cells have a period where the probability of quiescence fluctuates around 0.5. However, there was a distinct point after which the cell fate probabilities consistently pointed to one fate over another (Figure 6D). We have observed a similar point for the decision to undergo meiosis or not in our previous work (Argüello-Miranda et al., 2018) and we defined this point as the ‘decision point’ (DP). We define the DP as the time point after which all predictions were correct for a given cell, or equivalently after which the correct cell fate probability was always above 0.5.

We found that the average DP is 3.6 hrs after the initiation of AGR (Figure 6E). Note that the average probability of quiescence is about 0.5 before DP for both quiescent and senescent cells (0.52±0.1 /0.52±0.1 Quiescent/Senescent, mean±SEM). After the DP, the probability for quiescent cells became almost 1 for quiescent cells, and close to 0 for senescent cells (0.93±0.05 / 0.10±0.05 Quiescent/Senescent, mean±SEM). Also, like the percentage of correctly predicted cells, the DP depends on the biomarkers used to calculate the cell fate probabilities. Thus, based on the biomarkers we have examined in this study, our data suggests that the majority of cells “decide” on a fate within 3-4 hours of the initiation of the AGR phase, which is 6-7 hours before glucose replenishment and the manifestation of those cell fates.

## Discussion

In the field of cellular decision-making most research has been devoted to cell fate-specific signaling (Balázsi, van Oudenaarden, & Collins, 2011), such as the cell-cycle machinery (Doncic, Falleur-Fettig, & Skotheim, 2011). How general cellular homeostasis and organelle physiology affect fate decisions remains underexplored yet represent some of the most critical aspects of adaptive metabolism that influence cell fitness and survival. Here, we report that in response to acute glucose removal, isogenic yeast manifest distinct quiescent and senescent cell fates that correlate with temporally unique signature profiles (Figure 7). These include biomarkers associated with stress response transcriptional changes, interorganelle contacts, and cellular metabolism. Notably, these biomarkers display diverging signal profiles throughout AGR stress. We also identify an early predictor biomarker, Rim15, whose cellular abundance prior to AGR is highly correlated to post-AGR cell fate. Thus, our work pioneers a shift in focus from cell fate-specific signaling to other noncanonical factors related to nutrient signaling, interorganelle communication, stress response, and metabolism.

**Figure 7:**
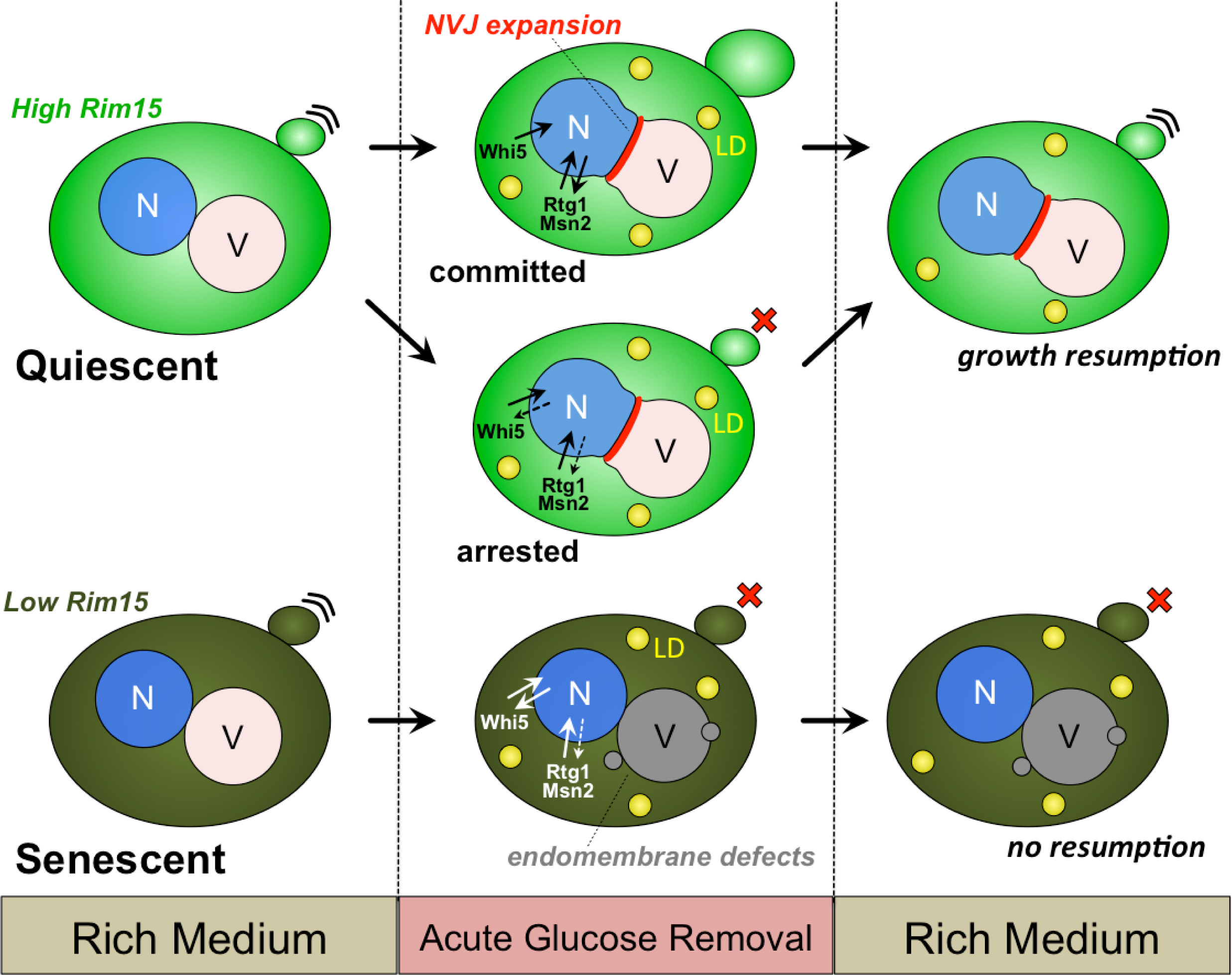
Working model of non-canonical determinants in cell fate.

In (Argüello-Miranda et al., 2018; Doncic et al., 2013; Wood & Doncic, 2019) we developed an experimental framework that merges microfluidics with live-cell imaging, allowing for simultaneous measurement of up to six-fluorescent channels, and an analysis pipeline that exploits algorithms for automated segmentation and cell tracking as well as automated extraction of single-cell data. Here we employed this powerful framework to the quiescence/senescence decision of budding yeast, allowing us to track single cells before, during, and after acute glucose removal while simultaneously monitoring multiple biomarkers. In (Argüello-Miranda et al., 2018) we combined our single-cell pipeline with a theoretical framework for forecasting cell fates and accurately predicted the fates of individual cells before the activation of cell fate-specific signaling in the context of cells undergoing meiosis. Here, we accurately predict cell fates in quiescence/senescence decision-making using this framework. In particular, we show that the quiescence/senescence decision is made before return to glucose-rich medium. Importantly, here and in (Argüello-Miranda et al., 2018) we propose that cells manifest a ‘decision point’ (DP), which marks the establishment of the predisposition for a cell fate, but does not mark an irreversible commitment.

Our work also adds to the temporal understanding of the cell-cycle marker Whi5 and its relationship to acute glucose loss. We show that even cells that pass Start will halt or postpone their cell cycle progression upon AGR. We also show that Whi5 accumulates in the nucleus in higher amounts in AGR stress compared to nutrient-rich media. Also, in a majority of these post-Start cells which translocated Whi5 out of their nuclei before AGR, Whi5 translocates back into the nucleus during sustained AGR, thus indicating a return-to-G1 phenotype. This suggests that Whi5 translocation out of the nucleus does not simply mark the commitment to the cell cycle in response to AGR. Notably, this surprising behavior of Whi5 has been independently observed by at least two other labs [Irvali et al., in preparation as communicated by JC Ewald; (Qu et al., 2019)].

Glucose starvation is known to drive interorganelle crosstalk and the expansion of the NVJ that connects the ER network to the yeast vacuole. Surprisingly we show that the NVJ tether Nvj1 is a highly predictive biomarker for cell fate following AGR stress. Nearly all quiescent cells expand their NVJ contact site during AGR, whereas senescent cells fail to do this. To our knowledge, this study is the first to show an interorganelle contact tether to be predictive of a cellular decision-making process. Given that NVJ contacts play important roles in lipid metabolism and autophagy (Roberts et al., 2003), this may indicate that NVJ-related autophagy is important for quiescence. In addition, interorganelle contacts like the NVJ enable collaboration and lipid exchange between different organelles, and thus execution of metabolic responses upon various environmental cues (Hariri et al., 2018). However, it should be noted that *nvj1*Δ yeast displayed similar cell fate profiles as wildtype yeast, indicating that although increased Nvj1 signal during AGR correlates with quiescence, Nvj1 loss does not alter the ability of cells to become quiescent.

LDs are key nutrient reservoirs, and studies indicate senescent cells increase their LD stores (Flor et al., 2017). However, a major knowledge gap is whether LDs are required for senescence, or merely a byproduct of its manifestation. Here we resolve this controversy in yeast by examining the LD marker Erg6 in post-AGR cell fate. As expected, AGR induced an increase in Erg6 signal as well as an increase in total neutral lipids, consistent with elevated lipid storage following glucose restriction. Surprisingly, LDs were not required for senescence or other cell fates, as yeast lacking the TG lipases (*tgl3,4,5*ΔΔΔ) and yeast entirely devoid of LDs (ΔLD) displayed similar proportions of quiescent cells, and only slightly reduced numbers of senescent cells. Given that LDs are storage organelles that enable cell survival during long-term (i.e. multi-day scale) starvation, this may indicate that LDs are dispensable for relatively short-term (i.e. multi-hour) adaptations to acute nutrient shortages.

Since senescent cells failed to resume budding upon glucose replenishment, we interrogated whether they could respond to external environmental cues and focused on their ability to internalize and deliver Mup1 to the vacuole. Although the majority of the senescent cells endocytosed Mup1, about half of the ones that endocytosed Mup1 did not successfully deliver it into the vacuole lumen for degradation. This protein sorting defect may be due to altered endomembrane trafficking and/or perturbed vacuole homeostasis, and in support of this senescent cells display altered CoV signatures of Vma1-mNeptune2.5 during AGR stress. In line with this, an intact vacuole is necessary for cell cycle progression (Jin & Weisman, 2015). Also, the vacuole-localized TORC1-Sch9 pathway can only signal from a mature functional vacuole, which may be compromised in senescent cells. Organelle homeostasis has been linked to cell fate responses in starvation, as mitochondrial morphology was found to be predictive whether a cell will proliferate or not after starvation upon glucose re-introduction (Bagamery, Justman, Garner, & Murray, 2020; Laporte et al., 2018). These collective observations indicate that organelle function is tightly connected to cell cycle progression, and to cell fate determination.

In an effort to identify a biomarker that provided predictive capacity to cell fate early in our experimental protocol, our work identified Rim15, a naturally low-abundance nutrient signaling kinase whose signal prior to AGR can accurately predict post-AGR cell fate. This supports a model where natural variation in the abundance of a nutrient-sensing effector can be important in the context of the acute nutrient stress, since cells must initiate fast responses to acute nutrient loss with existing protein machinery. This also indicates that long-term monitoring of single cells enables us to follow cell-cell differences back to upstream factors of cell commitment and that stochastic effects prior to starvation can affect long-term cellular responses.

In conclusion, our work uncovers new early predictors of cell fate, which surprisingly function in diverse cellular pathways ranging from stress response, to interorganelle communication, to endomembrane trafficking. These factors largely represent noncanonical determinants of cell fate in that they are not part of the cell cycle machinery. However, it should be noted that although these factors correlate with specific cell fates, their loss generally does not impact cell fate decision-making *per se*. This is exemplified by Nvj1, whose elevated signal strongly correlates with quiescence, even though *nvj1*Δ cells still undergo quiescence following AGR. Despite this, their predictive power highlights that, in response to acute nutrient shortage, specific signaling pathways are tightly integrated with organelle homeostasis, metabolism, and ultimately cell decision making.

## Acknowledgements

We sincerely thank to Sandra Schmid, Jonathan Friedman, Milo Lin, and Jennifer Ewald for critical feedback on the manuscript. We also thank Dr. Yanjie Liu, Sean Rogers, Natalie Ortiz, Dr. Orlando Argüello-Miranda, and Dr. Jungsik Noh for reagents and technical assistance. We also acknowledge our friend and departed colleague Dr. Andreas Doncic, whose scientific achievements served as a foundation for this work.

## Author Contributions

N Ezgi Wood, Conceptualization, Methodology, Investigation, Data Curation, Software, Formal Analysis, Validation, Writing, Visualization. Piya Kositangool, Strain Construction, Preliminary Data, Formal Analysis. Hanaa Hariri, TLC Experiments. Feedback on data interpretation and writing. Ashley Marchand, Strain Construction. Mike Henne, Conceptualization, Methodology, Writing, Supervision, Funding Acquisition.

## Figure Legends

**Figure S1**: **An overview of strains and their fluorescent biomarkers**.

**A)** An overview of the biomarkers that are discussed throughout the manuscript. **B)** Growth of the WT strains on the YPD plate, compared to the isogenic blank, which was on the same plate. **C)** Example image of 5-color imaging at the beginning and at the end of AGR. This colony is also shown in Video 1 and 3. **D)** Example quiescence-committed cell which displays “back-to-G1” phenotype. The mother cell (orange) arrests with a bud (dark blue) and has Whi5 already translocated out of the nucleus prior to bud emergence. During regular cell cycle progression, Whi5 should enter to the nuclei of mother and the daughter at the same time after completion of the mitosis. However, upon glucose removal, Whi5 translocates to the mother’s nucleus for about 5 hours, then leaves it. Later, it enters the nuclei of the mother and the daughter at the same time. **E)** Right, dead vs senescent cells during AGR. Note that dead cells usually have autofluorescence in multiple channels during the AGR assay as shown in the example. In contrast, senescent cells do not have autofluorescence, and have Vma1-mNeptune2.5 signal that shows an intact vacuole and nuclear Msn2-mNeonGreen localization. Left, dead cells during replenishment. Some dead cells do not autofluoresce, however, their phase image under rich conditions show distorted cell boundaries and the biomarkers do not reflect intact cellular structures. **F)** Nuclear Whi5-mKoκ traces, where WT-MSN2 and WT-RTG1 strains are shown separately. Cells are first ordered by their cell fate, and then ordered according to their Whi5-mKoκ traces during AGR.

**Figure S2: Analysis of biomarkers and endocytic uptake in cell fate populations.**

**A**,**B)** Comparison of nuclear Whi5-mKoκ and nuclear Msn2-/Rtg1-mNeongreen signals. Quiescent-committed cells translocate Msn2/Rtg1 out of the nucleus before they complete their cell cycle. Right, the traces of an example cell. The example cell translocates Msn2/Rtg1 to the nucleus upon glucose removal (orange lines). After a while, that signal decreases and the cell completes its cell cycle as indicated by the Whi5 translocation to the nucleus (black lines). Left, average nuclear intensities for Msn2/Rtg1 (orange) and Whi5 (black) for quiescence-committed cells (mean±SEM). Note that the nuclear translocation is not synchronized among cells, thus, averaging smoothens the curves. The simultaneous Whi5 translocation to the mother and bud’s nucleus is always sharp like in the example cells shown. **C)** Percentage of cells that endocytosed Mup1 and delivered it to the vacuole by cell fate during methionine responsiveness assay with SC-Methionine. **D)** Vma1-mNeptune average coefficient of variation (CoV) for quiescent (brown) and senescent (blue) cells (mean±SEM), which has been shown to be a measure of relative V-ATPase assembly (Dechant et al., 2010). The CoV normalized to 1 at the beginning of AGR decreases to 0.655±0.011 for WT-MSN2 quiescent cells and to 0.657±0.011 for WT-RTG1 quiescent cells, whereas it decreases to 0.558±0.012 for WT-MSN2 senescent cells and to 0.630±0.016 for WT-RTG1 senescent cells (mean±SEM). **E)** Average Vph1-GFP CoV for quiescent (brown) and senescent cells (blue) (N=26/10 Quiescent/Senescent).

**Figure S3: Analysis of NVJ expansion.**

**A**,**B)** NVJ expansion for WT-MSN2 and WT-RTG1, plotted separately. Same as Figure 3 B, but the quiescent cell population is reported by dividing them into quiescent-committed (orange) and quiescent-arrested (brown) cells.

**Figure S4**: **Analysis of Erg6-mTFP1 signal.**

**A) A, B)** Erg6-mTFP1 quantification as in Figure 4B, divided into three fates, strains shown separately. **C**,**D)** Erg6-mTFP1 mean intensity fold change normalized to the beginning of AGR. Averages are mean±SEM. **E)** Cell fate percentages by the strain, quiescent cells are divided into committed and arrested populations.

**Figure S5: Rim15 signal is distinct in cell fate populations.**

**A)** Rim15-mNeongreen signal quantification by three fates and the blank. Note that all cells including senescent cells with low Rim15 has higher Rim15 signal than the blank cells in that channel, indicating that low Rim15 signal is not background.

**Figure S6: Analysis of multi-parametric predictions.**

**A)** Percentage of correctly predicted cells using single parameters and the combination of parameters. Bold dashed line at 50% marks random guess. **B)** Percentage of correctly predicted cells using cell size. **C)** Same as Figure 6C, divided by strain. **D)** Average quiescence fate probabilities for quiescent (brown) and senescent (blue) cells calculated using single parameters. Compare these to Rim15 in Figure 6C. The combination of these parameters is used for calculating the quiescence fate probability in Figure S6C. **E)** Percentage of correctly predicted cells using Rim15-mNeonGreen mean intensity. Bold dashed line at 50% marks random guess. **F)** Percentage of correctly predicted cells, when predictions are done on cells which have signal one standard deviation above or below the population mean, using Rim15-mNeonGreen (red), Nvj1 mean intensity (blue), or Nvj1 area (yellow). **G)** Same as Figure 6E, divided by strain.

## Videos Legends

**Video 1: Acute glucose restriction (AGR) assay and fates.** The cells are marked according to their fates: quiescent-committed (orange), quiescent-arrested (green), and senescent (blue). The bottom quiescent-committed cell is recognizable by bud-growth during AGR. The top quiescent-committed cell is determined by simultaneous Whi5-mKoκ translocation to nuclei of mother and daughter during AGR (See Video3). Quiescent-arrested cell is scored by its bud growth and new bud emergence during glucose replenishment. Senescent cells did not resume mitotic cycles until the end of our assay.

**Video 2: AGR assay and 5-color imaging with WT-MSN2.** The cells are marked according to their fates: quiescent-committed (orange), quiescent-arrested (green), and senescent (blue).

**Video 3: AGR assay and 5-color imaging with WT-RTG1.** The cells are marked according to their fates: quiescent-committed (orange), quiescent-arrested (green), and senescent (blue). The phase image of this colony is also shown in Video 1.

**Video 4: Mup1 responsiveness assay with SC+Met.** Cells are marked according to their fates: quiescent-arrested (green) and senescent (blue). The bottom two channels show the superimposition of Vma1-mNeptune2.5 (purple) and Mup1-mKoκ/-pHluorin (green) channels to locate the Mup1 signal with respect to vacuole.

**Video 5: Mup1 responsiveness assay with SC+Met**. Cells are marked according to their fates: quiescent-committed (orange) and quiescent-arrested (green). The bottom two channels show the superimposition of Vma1-mNeptune2.5 (purple) and Mup1-mKoκ/-pHluorin (green) channels to locate the Mup1 signal with respect to vacuole.

**Video 6: Mup1 responsiveness assay with SC-Met.** Cells are marked according to their fates: quiescent-arrested (green) and senescent (blue). The bottom two channels show the superimposition of Vma1-mNeptune2.5 (purple) and Mup1-mKoκ/-pHluorin (green) channels to locate the Mup1 signal with respect to vacuole.

**Video 7: AGR assay and 5-color imaging with WT-RIM15.** The senescent cells are marked blue. The rest of the cells are quiescent-arrested.

