## Supplementary figures and images for "Nutrient signaling, stress response, and interorganelle communication are non-canonical determinants of cell fate"

### Figure S1

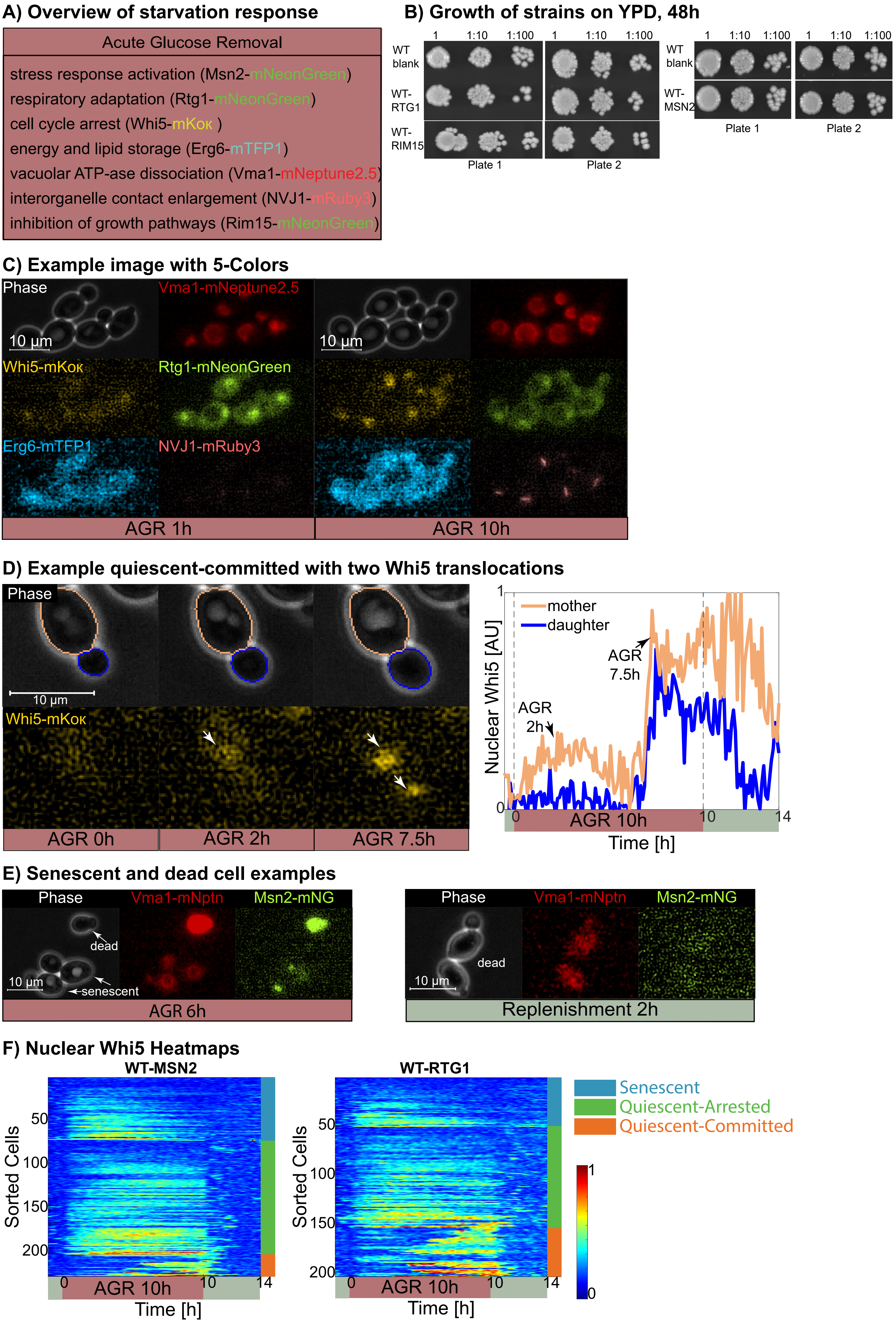

### Figure S2

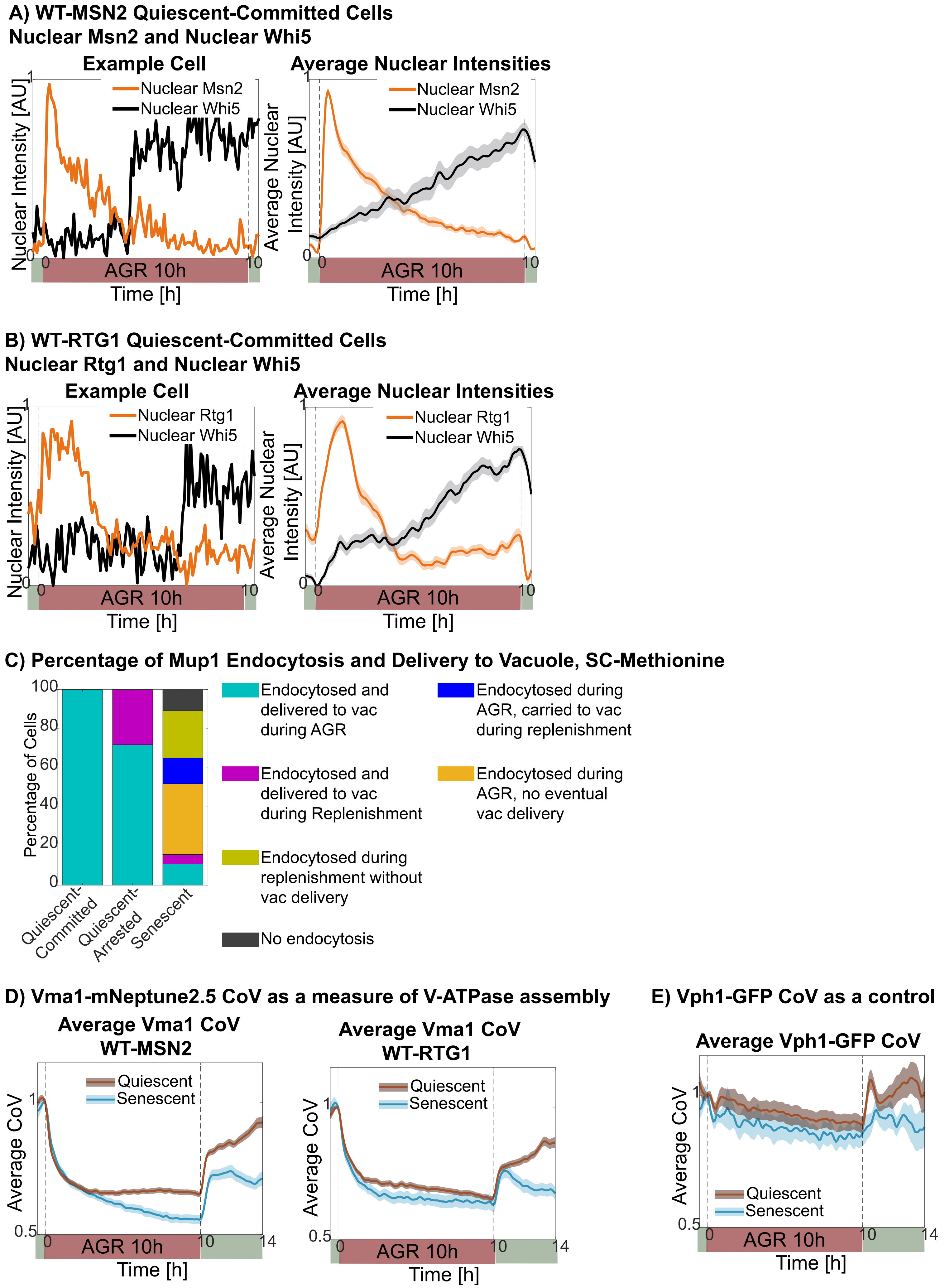

### Figure S3

**A) NVJ Expansion WT-MSN2**

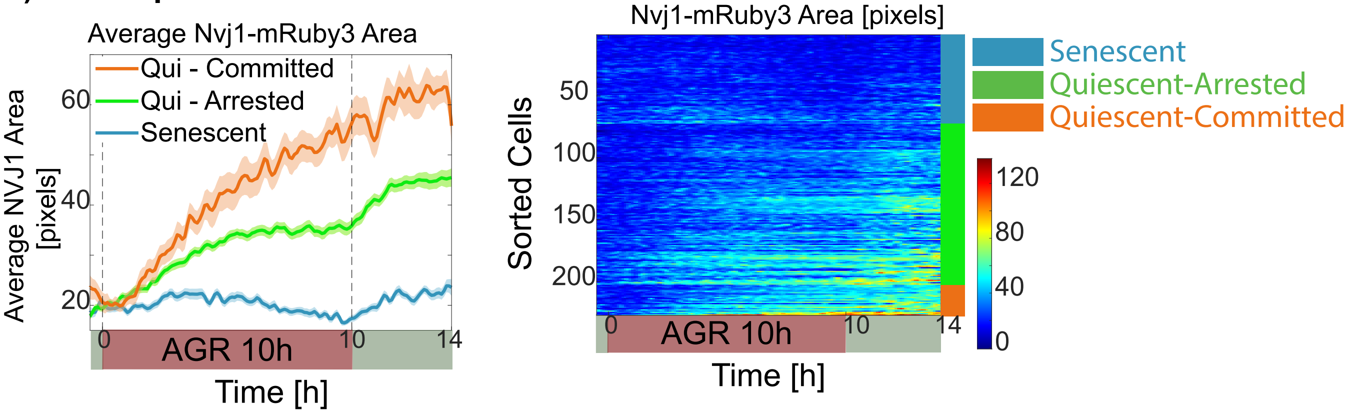


**B) NVJ Expansion WT-RTG1**

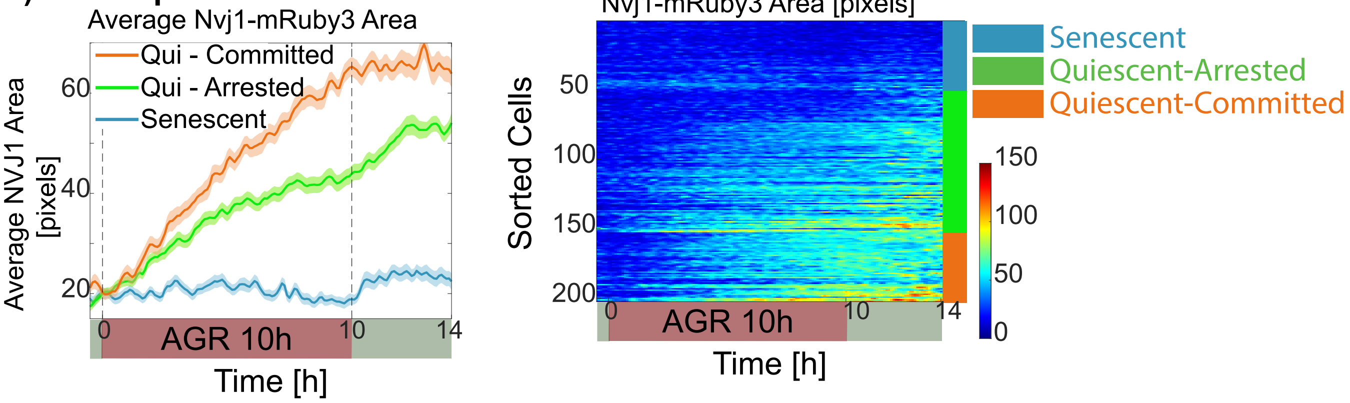

### Figure S5

#### A) Rim15-mNeonGreen Signal WT-RIM15

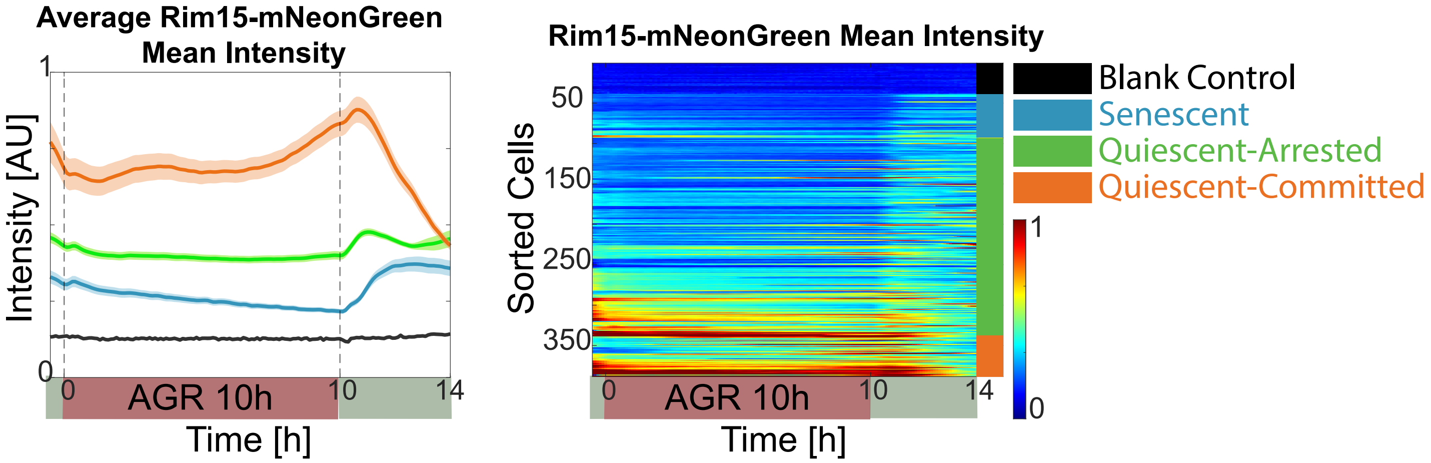

### Figure S6

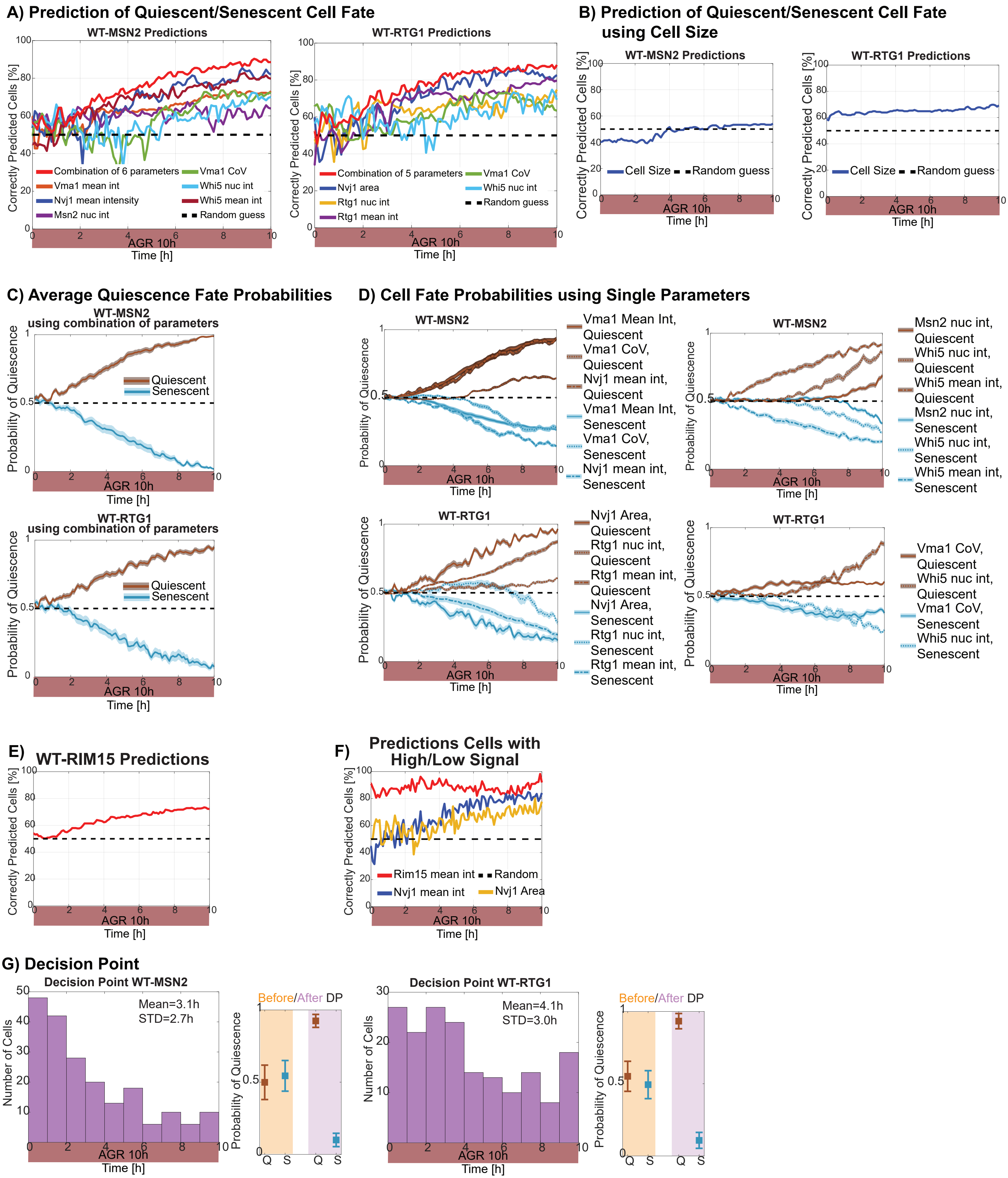
