## Supplementary material for "Nutrient signaling, stress response, and interorganelle communication are non-canonical determinants of cell fate": Figure S4

A) Erg6-mTFP1 Signal WT-MSN2

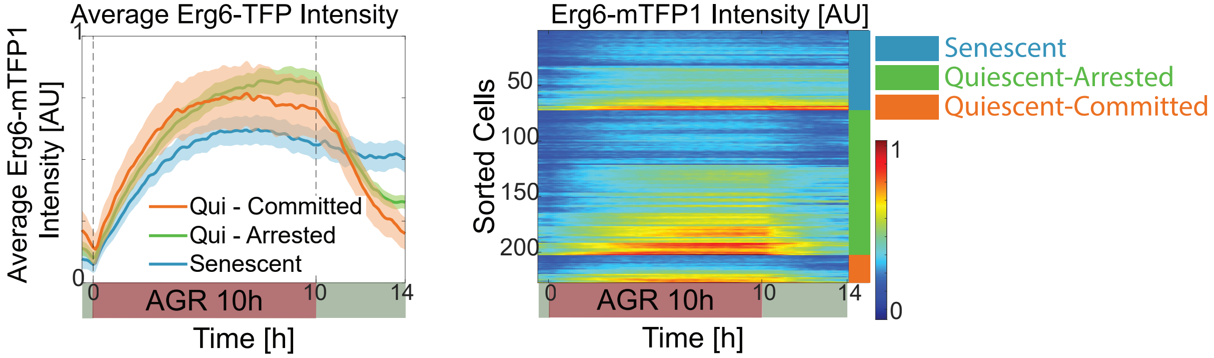


B) Erg6-mTFP1 Signal WT-RTG1

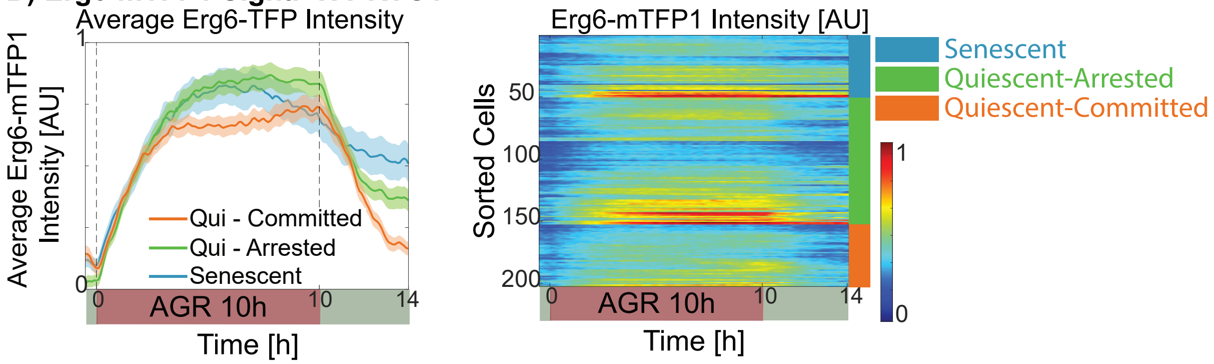


C) Erg6-mTFP1 Signal Fold Change WT-MSN2

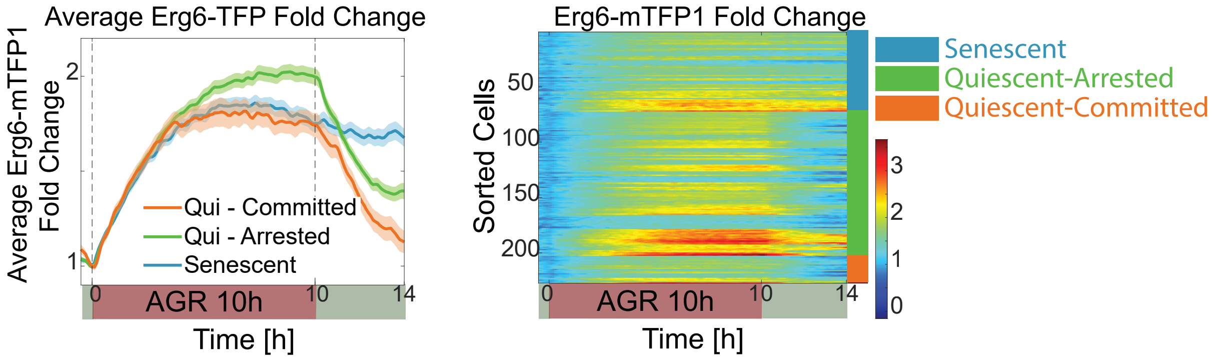


D) Erg6-mTFP1 Signal Fold Change WT-RTG1

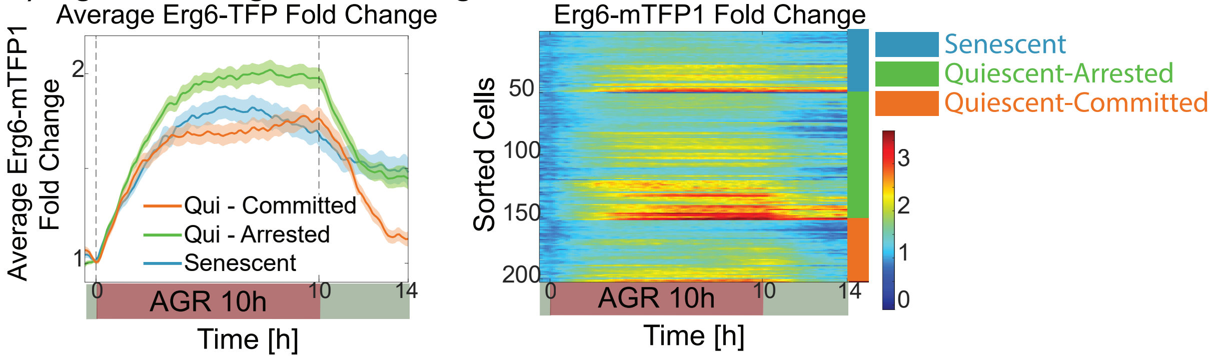


E) Cell Fate Percentages by Strain

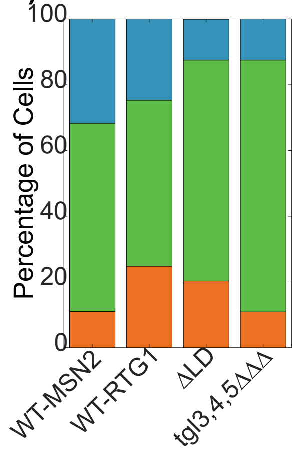
